# High-quality chromosome-level genomes of two tilapia species reveal their evolution of repeat sequences and sex chromosomes

**DOI:** 10.1101/2020.03.30.016618

**Authors:** Wenjing Tao, Luohao Xu, Lin Zhao, Zexian Zhu, Xin Wu, Qianwen Min, Deshou Wang, Qi Zhou

## Abstract

**Background:** Tilapias are one of the most farmed fishes that are coined as ‘aquatic chicken’ by the food industry. Like many other teleosts, Nile tilapia and blue tilapia exhibit very recent transition of sex chromosome systems since their divergence about 5 million years ago, making them a great model for elucidating the molecular and evolutionary mechanisms of sex chromosome turnovers. Studies into their sex-determining pathways are also critical for developing genetic sex control in aquaculture.

**Results:** We report here the newly produced genomes of Nile tilapia and blue tilapia that integrate long-read sequencing and chromatin conformation data. The two nearly complete genomes have anchored over 97% of the sequences into linkage groups (LGs), and assembled majorities of complex repetitive regions including telomeres, centromeres and rDNA clusters. In particular, we inferred two episodes of repeat expansion at LG3 respectively in the ancestor of cichlids and that of tilapias. The consequential large heterochromatic region concentrated at one end of LG3 comprises tandem arrays of mRNA and small RNA genes, among which we have identified a candidate female determining gene *Paics* in blue tilapia. *Paics* show female-specific patterns of single-nucleotide variants, copy numbers and expression patterns in gonads during early gonadogenesis.

**Conclusions:** Our work provide a very important genomic resource for functional studies of cichlids, and suggested that unequal distribution of repeat content that impacts the local recombination rate might make some chromosomes more likely to become sex chromosomes.

## Introduction

Tilapias belong to the largest vertebrate family of African cichlids (about 3000 species, order Perciformes) that underwent explosive speciation within the last 10 million years (MY) [1-3]. While two thirds of the cichlids are mainly endemic in lakes of East Africa, and are used as the textbook model for studying mechanisms of sympatric speciation; various tilapia species successfully colonized a much wider range of habitats and have become some of the most important aquaculture species. In particular, the earliest record of raising Nile tilapia (*Oreochromis niloticus, ON*) can be dated back to Ancient Egypt. Now it is projected to soon overtake carp and salmon as the most important farmed fish. A second popular tilapia species, blue tilapia (*Oreochromis aureus, OA*) diverged from *ON* less than 5 MY ago [4], and has a better cold and saline tolerance, thus is frequently used to produce hybrids with *ON*. Tilapia species from the *Oreochromis* and *Sarotherodon* genera, and many East African cichlids are mouthbrooders [5]. That is, females undergo periods of fasting when brooding the eggs, and sometimes even caring for the fry for extended time. Such a tremendous energy cost of females is one of the major causes that render the larger-sized males the favored sex in tilapia aquaculture. The current predominant practice of sex control in tilapia production is to use the cost-effective hormones rather than adjusting the temperature or population density to induce sex reversal, despite the potential risks to the consumers and the environment [6, 7]. This is mainly due to the lack of detailed knowledge about the genetic sex determining (GSD) pathways of tilapia species.

There is a strong and persistent interest in studying the tilapia SD mechanisms and sex chromosomes, in order to produce all-male fingerlings, and also to use tilapias as a model to unravel the molecular and evolutionary mechanisms of vertebrate sex chromosome turnovers [8-10]. In contrast to the conserved and stable sex chromosomes within mammals, birds or Drosophila, teleost fish harbor a remarkable diversity of male heterogametic (XY, like that of mammals), female heterogametic (ZW, like that of birds), and environmental SD (ESD) mechanisms frequently between sister species [11-13]. Fish sex chromosomes also do not usually exhibit a high degree of differentiation [13-15], which hampers the identification of the sex chromosomes or the exact SD region cytologically. Some species like *ON* combine both GSD and ESD, suggesting sex in these species is a threshold trait that can be determined by genetic and environmental factors [9]. Despite the complexity of SD systems, and a lack of abundant genomic resources and functional genetic tools until very recently, there have been great efforts of mapping the SD regions among the tilapia species. Early inspection of synaptonemal complex speculated that a large pair of chromosomes corresponding to linkage group 3 (LG3) with incomplete pairing at its terminals maybe the XY chromosome pair of *ON* [16-19]. However, genetic mapping using various types of markers (e.g., microsatellites) indicated that another chromosome LG1 carries an unknown male SD gene as an XY system [20, 21]. The SD region was recently narrowed down into a 9Mb region, through mapping the Illumina reads of both sexes against a high-quality LG1 sequence generated by PacBio reads from a female Egyptian strain of *ON* (*ONEg*). Similarly, by mapping the reads of *OA*, the SD region was inferred to span 50Mb of LG3, as a ZW system [22]. The rapid transition of sex chromosome system between the two species *OA* and *ON* occurred within only 5 MY. More strikingly, another study has mapped the male SD gene in a Japanese strain of *ON* (*ONJp*) onto LG23 rather than LG1 [23-26]. The Y-linked male SD gene is a duplicated copy of *anti-Mullerian hormone* (*Amhy*), and its disruption by CRISPR/Cas9 causes male-to-female sex reversal [25]. This is the first functionally validated SD gene of cichlids, and has demonstrated a probably even more recent turnover of SD genes between tilapias.

We present here the chromosome-level genome assemblies and comparative analyses of *ONJp* and *OA* vs. the Lake Malawi cichlid species *Metriaclima zebra* (*MZ*). Besides their aquaculture significance, *ON* has been used as the outgroup for studying genomic mechanisms of cichlid species radiation [3]. Moreover, *ONJp* is the first cichlid stock on which transgenics and gene-editing have been successfully conducted [25, 27, 28], with the demonstrated potentials for future functional studies of cichlids. We harnessed the single-molecule real-time sequencing technology and produced highly accurate and continuous assemblies of the homogametic sexes of *ONJp* and *OA*, covering their rDNA clusters and centromeric regions. By incorporating chromatin conformation (Hi-C) data, we further anchored over 97% genome sequences of each species into linkage groups, in particular a large heterochromatic region of small RNA gene clusters located at the terminal of LG3. Finally, we narrowed down the SD regions of both species and provided insights into the history of sex chromosome turnover of both species.

## Results

### Genome assembly and annotation of tilapia genomes

We produced 96× and 85× genomic coverage of Nanopore long-read sequences, with a read N50 length of 26kb and 39kb for a female *ONJp* individual (with XX genotype) and a male *OA* individual (ZZ genotype) respectively. Such a high sequencing coverage has overcome the higher error rate of Nanopore reads than that of PacBio reads, and produced similar numbers of genome size, contig N50 length and genome completeness measurement (BUSCO score), compared to those of *ONEg* and *MZ* previously derived from PacBio reads (**Figure 1a-b, Table 1**) [29, 30] or other chromosome-level fish genomes (**Supplementary Fig. S1**). With the linkage information provided by the Hi-C technology, we anchored 97.4% and 97.8% of the genome of *ONJp* and *OA* into chromosomes (**Supplementary Fig. S2**), followed by genome polishing with high coverage of Illumina reads and manual curation of scaffold orders within the chromosomes. The percentages of anchored sequences of the two genomes are higher than that (90.2%) of *ONEg* by genetic map [30]. And notably, we found no interchromosomal and very few intrachromosomal rearrangements by the genome-wide comparison between *ONJp* and *ONEg*, confirming the correct orientation of scaffolds within our chromosome assemblies. The unanchored sequences are enriched for repetitive elements that have alignments with multiple anchored chromosomal sequences. The most significant improvement of *ONJp* over *ONEg* is concentrated at the highly repetitive end of LG3, which made LG3 the largest assembled chromosome (over 130 Mb) in the genome (**Figure 1c, Supplementary Table S1**). Its overall repeat content (63%) is estimated to be around 2 fold higher than any other chromosomes in the genome (**Figure 1d**), and such a large chromosome-specific heterochromatic region has been found in both *ONJp* and *OA* (**Supplementary Fig. S2**). Particularly, the last 70 Mb sequence of LG3 exhibits an extremely high repeat content of about 75%. This is consistent with previous cytogenetic studies that identified LG3 as the largest characteristic subtelocentric chromosome shared by all the examined Tilapiine species [16, 31].

**Table 1.** Genome assembly and annotation statistics.

| **Assembly** | **ONEg** | **ONJp** | **OA** |
| --- | --- | --- | --- |
| Sequencing platform | Pacbio | Nanopore | Nanopore |
| Coverage | 44X | 96X | 85X |
| Assembly size | 1,005,681,550 | 993,468,885 | 1,005,590,959 |
| Contig N50 | 2,923,640 | 2,651,554 | 4,404,323 |
| # contig | 3,010 | 1,201 | 805 |
| Scaffold N50 | 38,839,487 | 40,346,024 | 40,723,988 |
| # scaffold | 2,460 | 403 | 303 |
| % anchored | 90.20% | 97.40% | 97.80% |
| complete BUSCOs | 97.00% | 97.30% | 97.50% |
| Fragmented BUSCOs | 1.60% | 1.40% | 1.30% |
| Missing BUSCOs | 1.40% | 1.30% | 1.20% |
| # gene | 29,537 | 25,264 | 25,467 |
| Repeat content | 36.5 | 40.4 | 39.4 |

**Figure 1.**
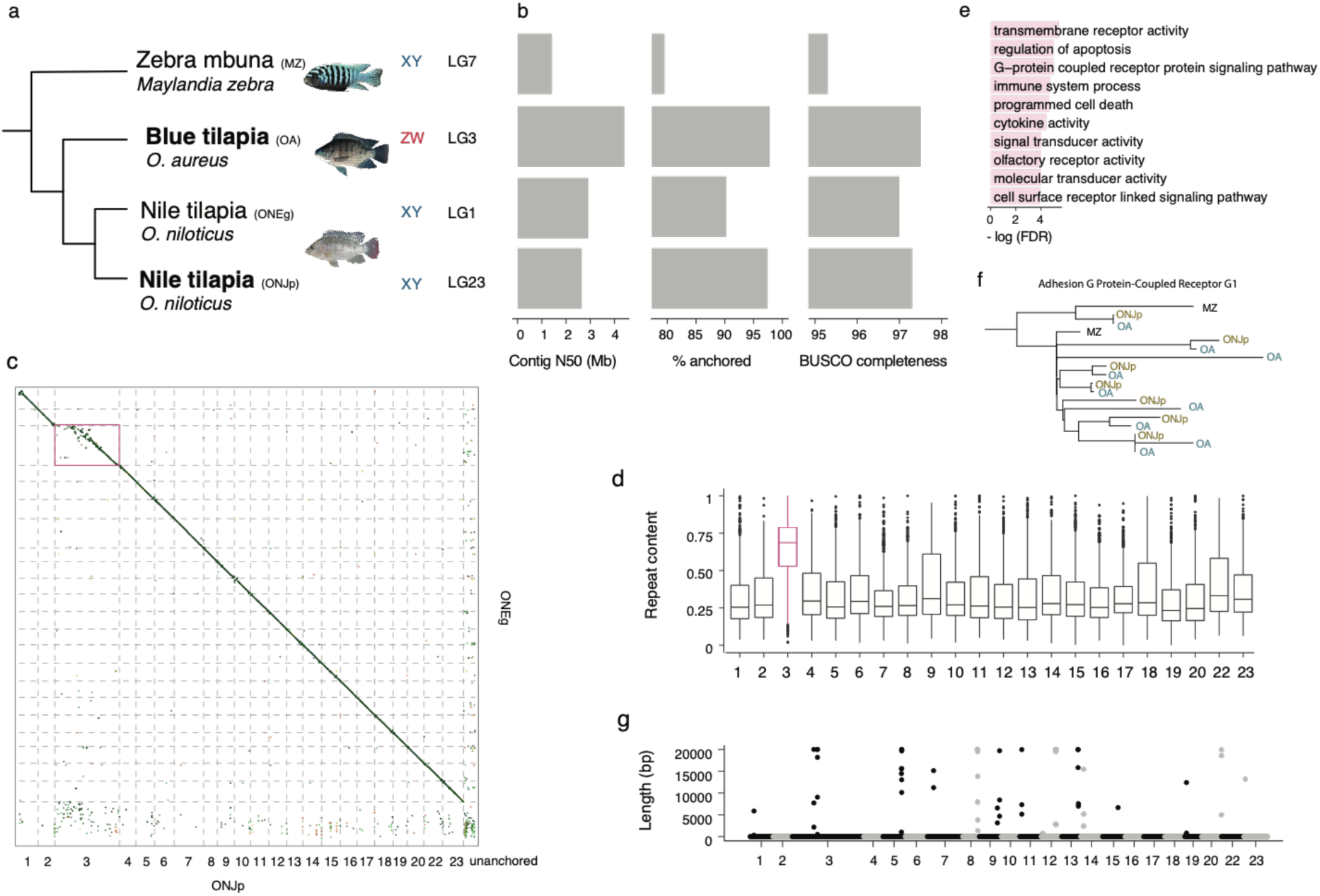
High-quality genome assemblies at the chromosome level. a) In this study, the genomes of the Japanese strain of Nile tilapia (*ONJp*) and blue tilapia (*OA*) were assembled. The genomes of the Egyptian strain of Nile tilapia (*ONEg*)[22] and Zebra mbuna (*MZ*)[30] have been published. The heterogamety (XY or ZW) and the sex chromosome (linkage group, or LG) were shown next to the fish photos. b) Contig N50, the percentage of sequences anchored into chromosome, and genome completeness (BUSCO) were compared among the four genomes. c) The genome synteny between *ONJp* and *ONEg* is highly conserved except for LG3 which is much larger in terms of anchored sequences for *ONJp* relative to *ONEg*. d) LG3 has a much higher repeat content than the other LGs. e) GO enrichment (FDR < 0.05) for gene families that have expanded in the tilapia lineage. Redundant GO terms were removed. f) One example of gene duplication at the ancestor of tilapias. g) The density of centromeric repeats (length per 50k) is shown along each chromosome.

The total number of annotated genes is comparable between the *MZ* cichlid versus the two tilapia species (**Table 1**). We found certain gene ontology (GO) categories of genes are enriched (FDR<0.05, **Supplementary Table S2**) for significant family expansion or contraction specifically at the ancestor of tilapias after their divergence from the other cichlids (**Supplementary Fig. S3**). For example, besides the reported olfactory receptor gene families [3, 32], we found immune-response related genes (e.g. *CTLA4*, **Figure 1e**), and ‘G-protein coupled receptor protein signaling pathway’ genes (**Figure 1f**) related to environmental sensing have specifically increased their copy numbers in the two tilapias. These genes may have contributed to tilapias’ adaptation to more varieties of ecological niches compared to the lake cichlids, and explained why they have been introduced as aquaculture species to over 150 countries.

### Characterization of complex repetitive genomic regions of tilapias

Non-coding repetitive sequences can play critical structural and regulatory roles in the genome, and there have been great efforts in mapping and characterizing such elements (e.g., satellites, short interspersed nuclear elements (SINE), rDNA) in the cichlids [33]; [34-37]. The two highly-continuous tilapia genomes allow us to scrutinize these highly repetitive genomic regions that are mostly absent or unanchored in the previous version of genome assemblies. There are two characteristic satellite sequences SATA and SATB present in large tandem arrays with up to hundreds of thousands of copies in the tilapia genomes. In particular, variants of SATA were previously identified to be concentrated at the centromeric regions of *ON* chromosomes [36, 38], and used as a phylogenetic marker to separate different tilapia tribes [37]. We used high (between 30 to 620) copy numbers of SATA as a marker and annotated the putative centromeric regions of over half of the chromosomes in both *ONJp* and *OA* genomes (**Figure 1g**). As expected, we found the locations of putative centromeres are colocalized with the junctions between the two arms of large intrachromosomal interaction domains (**Supplementary Fig. S4, Supplementary Table S3**), similar to what has been reported in other vertebrates [39]. The monomer sequence of SATA satellites shows high degrees of variations (indels or SNPs) at certain monomer positions between copies of the same or different chromosomes, but there are intriguingly no variations at all in the last 58bp region across all the mapped loci of the two species (**Supplementary Fig. S5**). This suggests concerted evolution and potential functional constraints within this region. Most assembled putative centromeres are close to or at the tip of the chromosomes. Their genomic locations are conserved between the *ON* and *OA* genomes, without obvious centromere repositioning events that may play a role in speciation [40] (**Supplementary Fig. S6**). This is in accordance with the reported highly conserved karyotype between blue and Nile tilapia species that consists of almost exclusively acrocentric or subtelocentric chromosomes [33, 41, 42]. The other satellite SATB of longer monomer length (1.9kb) [36] has been previously shown to be concentrated on the short arm of one chromosome, or on those of up to 14 pairs of chromosomes, depending on the experimental conditions of fluorescence *in situ* hybridization (FISH) [38]. We confirmed here that SATB satellite is frequently co-localized with SATA and enriched in pericentromeric regions of at least 8 chromosomes in both tilapias (**Supplementary Fig. S7**).

The other classic tandem array sequences that are of great interest but extremely difficult to assemble are the ribosomal DNA (rDNA) clusters with hundreds of thousands of copies in the genome. This is evidenced by the fact that rDNAs have been mapped for their locations in over 500 fish species, but are only studied for their partial genomic sequences in three species [43]. Eukaryotic rRNA genes are divided into two classes of 45S (corresponding to the nucleolar organizer regions, NORs) and 5S rRNA genes. They are transcribed by different RNA polymerases, and often located on different chromosomes in teleost species. Here we dissected the complex sequence structures and mapped the rRNA gene clusters in *ONJp* and *OA* genomes. Both species show similar numbers and chromosomal locations of mapped loci (**Supplementary Fig. S8**), but the *OA* genome (**Figure 2b**) probably captures a more complete sequence composition with its better assembly quality thus is used for demonstration here. We mapped the major 45S and 5S rRNA clusters respectively on LG14/6/4 and LG23/22, which is consistent with previous cytogenetic results [33]. The total copy numbers of 45S and 5S rDNA were estimated to be 123 and 171 throughout the *OA* genome. The 45S rDNA cluster on LG14 is located at the end of acrocentric chromosomes (**Figure 2a**), and consists of 11 transcriptional units coding for the 18S, 5.8S and 28S rRNAs. Each unit containing internal transcribed spacers (ITS) is separated by intergenic non-transcribed spacers (IGS). Three tandem units are organized in inverted orientation to the other eight units, suggesting recombination may happen between these units by forming a hairpin structure (**Figure 2c**). The IGSs of the two groups of tandem units are only partially homologous to each other in sequence, and are themselves tandem arrays of multimers. Remarkably, there are nine copies of 3.7 kb repetitive sequences (>97% sequence similarities between copies) in one IGS (**Figure 2c**), and each copy consists of 25 copies of 102bp sequences (about 95% sequence similarity between copies) that are separated into two clusters. Such nested tandem arrays of repetitive sequences resemble the higher order repeats (HORs) of human centromere [44] and remain to be studied for their functions.

**Figure 2.**
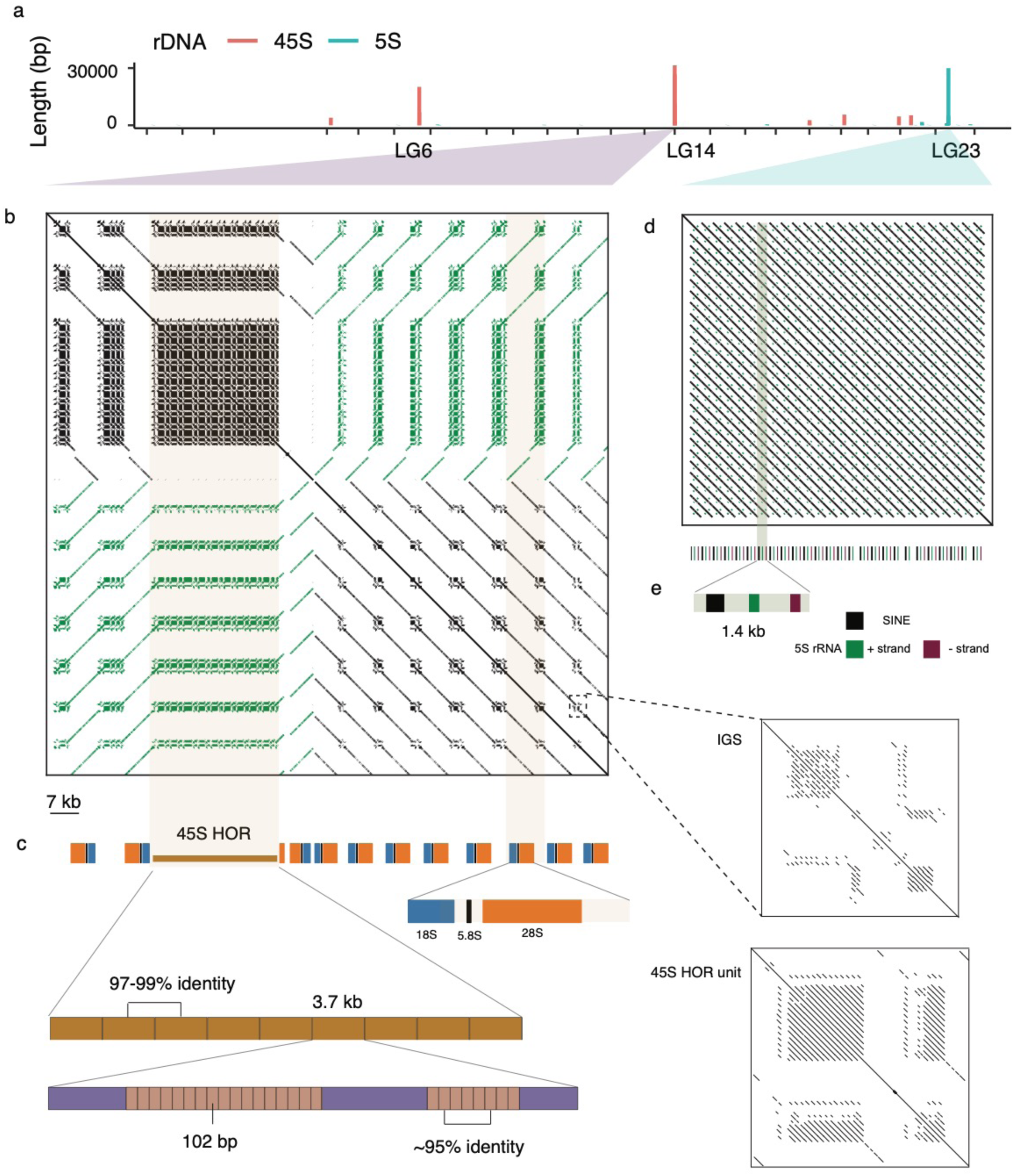
The genomic organization of rDNA loci. a) The length of rDNA sequence per 50kb along the chromosomes of *OA*. We selected one 45S locus (b) and one 5S locus (d) for demonstration. b) the dotplot showing an array of 11 copies of 45S genes and the intergenic spacers (IGSs). The green colors represent reversed alignments. One IGS was selected for a zoom-in view. c) Each 45S locus contains one 18S, one 5.8S and one 28S gene. The second IGS forms a higher order repeat (HOR), consisting of 9 repeats of a 3.7kb element which itself consists of tandem duplications of a 102 bp sequence. d) The dotplot showing an array of 25 copies of 5S rRNA gene loci. Each locus contains two 5S rRNAs with opposite coding directions and one SINE element shown in e).

There are two classes of 5S rDNA (**Figure 2e**), which respectively consist of 1.4kb (type I) and 0.5kb (type II) repeat units. The type I 5S rDNA cluster residing on the LG23 consists of 26 tandem duplications of repeat units. Each repeat unit contains a 5S RNA, an inverted 5S RNA and a SINE repeat. This SINE repeat seems to be derived from 5S RNA, with more than half of its sequence homologous to 5S RNA.

### LG3 heterochromatic regions encompass tandem arrays of protein-coding and small RNA genes

The heterochromatin concentrated at the end of LG3 forms a large unpaired region during male meiosis, which was presumed to be the sex chromosome pair of *ON* [18]. The repetitive nature of this part of LG3, together with its extremely low recombination, may have contributed to the difficulty of finding genetic markers to anchor a large portion of LG3 in the *ONEg* assembly [22]. Later QTL mapping and genomic analyses, however confirmed that LG1 or LG23 is the sex chromosome of *ON* [22, 26], while LG3 has been frequently adopted as the sex chromosome in other tilapia species [22, 45].

To trace the origin of such unusual autosome-specific heterochromatin, we compared the assembled sequences of LG3 of *ONJp* and *OA* vs. that of *MZ*. Based on their syntenic alignment (**Figure 3a**) and the distribution of repeat content along the LG3 (**Figure 3b**), we inferred that there were probably two episodes of repeat expansion (RE): the first one is shared by both *MZ* and *ONJp* (**Supplementary Fig. S9**), thus may have occurred at the ancestor of all cichlids. It is manifested as a turning point at around the 30Mb position, where the repeat content starts to increase, while the GC content and recombination rate start to decrease [30], forming a heterochromatic region (Het-1) with over 50% of the sequences as repetitive elements. The clear negative correlation between these genomic features can be explained by the scarcity of GC-biased gene conversion caused by the low recombination rate [46]. The second RE impacts the region beyond the 70Mb position (Het-2), and probably occurred more recently at the ancestor of tilapia species. It accounts for the dramatic size increase of LG3 from 45 Mb in *MZ* to more than 130 Mb in both tilapias, and the increase of repeat content from around 50% to over 70% (**Figure 3c**). This massive repeat expansion involves all repeat families but DNA transposons and simple repeats (**Figure 3b**). Most repeats seem to have started the expansion from the euchromatin region toward the other chromosome end (**Figure 3c**), the latter of which is enriched for the younger repeat elements that show a low level of sequence divergence from their consensus sequences (**Supplementary Fig S9**). Of particular interest are three previously uncharacterized repeat families: although they only account for 1.9% of all LG3 repeat sequences, they are almost exclusively concentrated on LG3, with two of them only at the LG3 Het-2 region (**Figure 3d, Supplementary Fig. S10a**). They are shared by all sequenced cichlids studied here (thus we named them as CLD repeats), and include one DNA transposon (DNA_CLD1), and two uncategorized repeats UNCLD1 and UNCLD2. But their copy numbers have specifically increased in tilapias: for other tilapia species without a genome generated by third-generation sequencing, we estimated their relative repeat copy numbers by kmer frequency scaled against the genome coverage, and found minor expansion of UNCLD1 and UNCLD2, but a 12.6 fold expansion of DNA_CLD1 across all sequenced tilapias relative to *MZ* (**Supplementary Fig S10b**). The bombardment of various repeats has clearly demarcated the entire chromosome of *OA* and *ONJp* into one large active (A) and one large repressive (B) chromatin compartment (**Figure 3e-f**), revealed by our chromatin interaction analyses. We artificially marked the boundary between the A/B compartment also as the one between Het-1/-2 regions. As expected, genes located at the Het-1 or -2 regions of LG3 are expressed at a significantly (*P* < 2.2e-16, Wilcoxon rank sum test) lower level than those on the other LGs across all examined tissues (**Figure 3g**).

**Figure 3.**
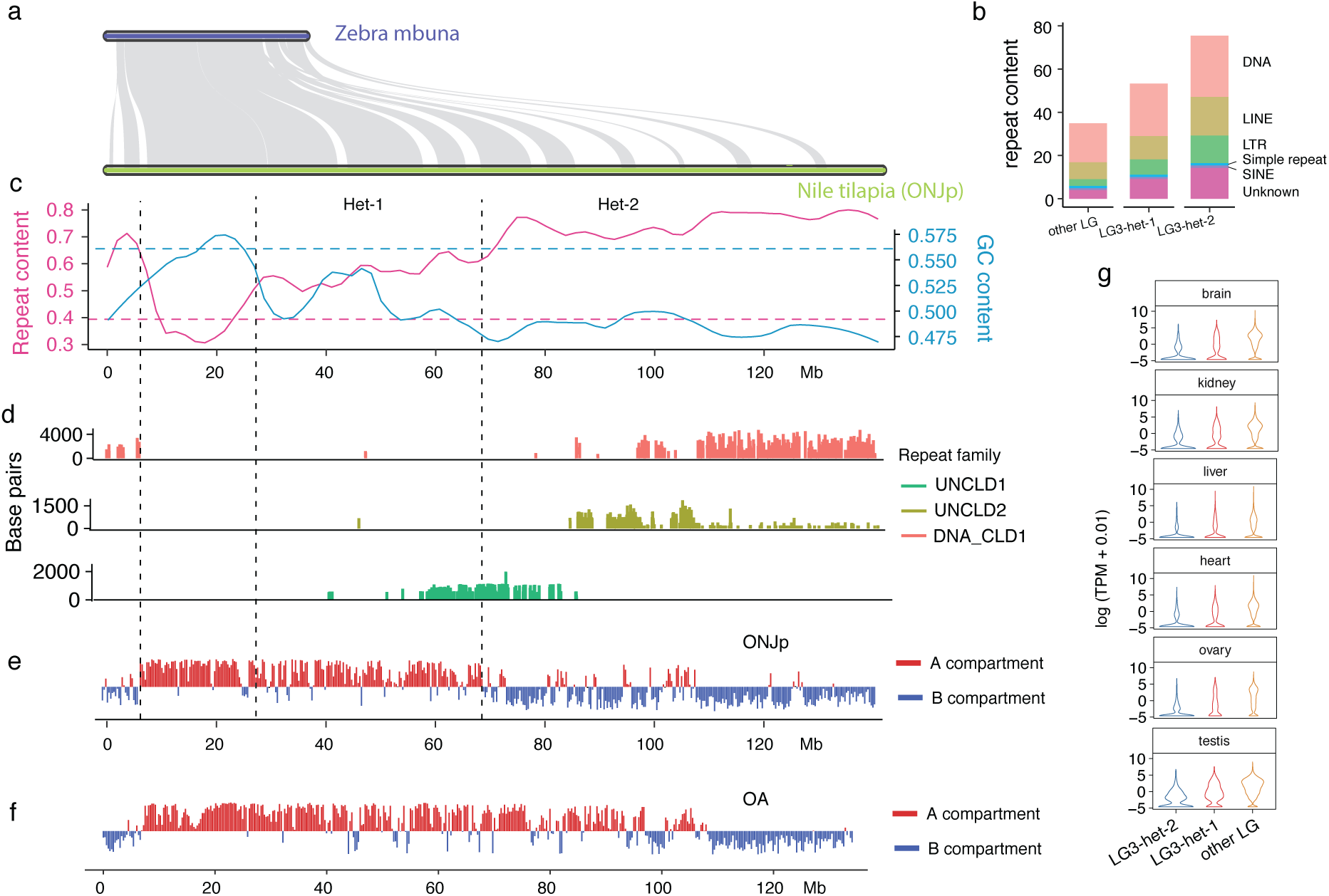
Heterochromatin region of LG3. a) The synteny between the LG3 of *MZ* and *ONJp*. Each grey band represents a synteny block. b) Comparison of the composition of repeats on two heterochromatic parts (Het-1 and Het-2) of LG3 and other LGs. c) The distribution of repeat content (pink) and GC content (blue) along the LG3 of *ONJp*. The heterochromatic part is divided into Het-1 and Het-2 which show differential degrees of heterochromatinization. d) The density (length per 5kb) of three transposable elements along the LG3. e-f) The distribution of A/B compartments on the LG3 of *OA* and *ONJp*. The A compartment usually corresponds to the active chromatin domain, and the B compartment corresponds to the repressive or heterochromatin domain. They are derived from eigenvector analyses of Hi-C data. g) Expression profiles of six tissues of *ONJp*. Both Het-1 and Het-2 regions have significantly lower expression levels compared with genes from other chromosomes.

The heterochromatic regions of LG3, however, are not gene deserts, but instead more frequently harbor gene duplications than other LGs (**Figure 4a**), probably due to the non-homologous recombination or replication slippage mediated by the excessive repeats [16]. This is exemplified by independently formed gene clusters of tandem duplication among the *MZ* and the two tilapia species within their Het-2 regions (**Figure 4b**). Some species-specific gene duplicates have probably evolved novel functions: for example, gene copies of *Zina33* are mainly expressed in the heart and kidney of *MZ*, but have acquired new expression patterns in brain and liver in *ONJp* in some copies (**Supplementary Fig. S11**). Besides facilitating the generation of these new gene duplicates that may contribute to the species-specific adaptation, we also found a disproportionately large number of predicted PIWI-interacting RNA (piRNA) or small-interfering RNA (siRNAs) encoding loci on LG3, which account for about 30% of the small RNA loci throughout the genome of *ONJp* (**Figure 4c**). Particularly, the predicted small RNA loci form a gradient along the LG3 of their density (number of loci per 100kb) and are mostly concentrated on the more recently formed Het-2 region (**Figure 4d-e**).

**Figure 4.**
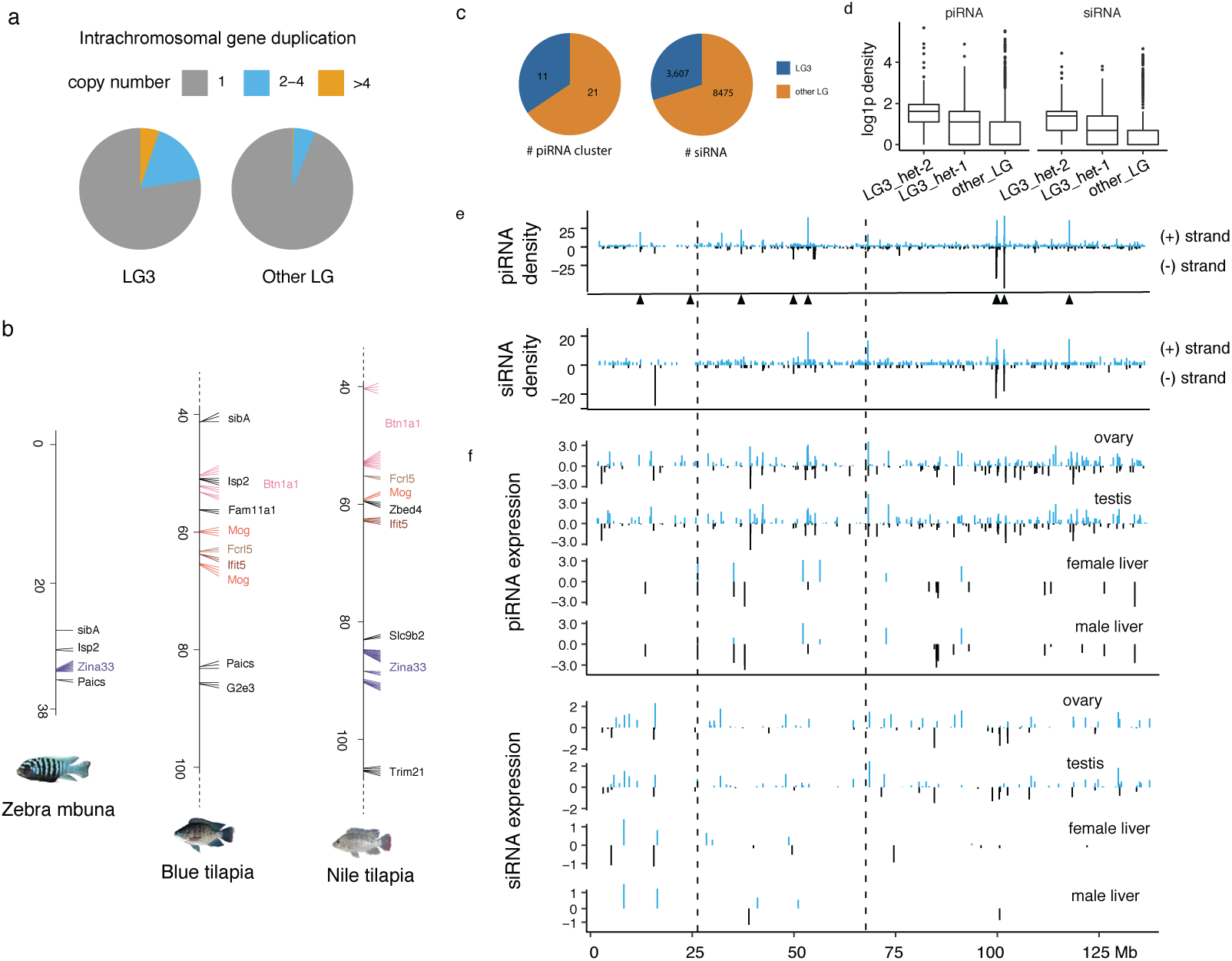
LG3 heterochromatin contains tandem arrays of mRNA and sRNA genes. a) LG3 has a larger portion of duplicated genes compared with other LGs. b) Tandem duplications of genes with at least two duplicated copies are shown along the LG3 of *OA* and *ONJp*. The homologous genes to tilapia duplicates are also shown on *MZ* LG3. Homologous genes of the same family are in the same color across species. For tilapias only the regions from 40 to 100 Mb of LG3 are shown. c) A disproportionately larger number of piRNA clusters and siRNA genes on the LG3. d) log1p transformed density of piRNA and siRNA genes over 100 kb windows. e) The density of piRNA and siRNA on the positive (blue) and negative (black) strand. The black triangles indicate the locations of piRNA clusters. f) log transformed expression levels (TPM) of piRNA and siRNA of gonads and livers on the positive (blue) and negative (black) strand. Genes with low expression (TPM < 1) were filtered out.

Similar to the reported expression patterns of piRNAs or siRNAs in other model species, these small RNAs are predominantly expressed in the gonads relative to the liver tissue (**Figure 4f**), suggesting they play a similar role of suppressing transposon activities and guard the germline genome integrity as they do in other species [47]. Interestingly, the piRNA-encoding repeat elements show a bimodal distribution of ages reflected by their sequence divergence level from the respective consensus sequences (**Supplementary Fig. S12**), with the peak of younger repeats largely overlapped with those of LG3. This together with the more concentrated distribution of small RNA loci at Het-2 provide evidence that the more recent RE of LG3 probably has selected for the emergence of novel small loci as a response to tame the new transpons acquired on LG3.

### Turnover of sex chromosomes and sex determination pathways

The complete X (LG23) chromosome sequence of *ONJp* and the Z chromosome (LG3) of *OA* provide us a great opportunity to gain insights into the evolution process and the consequences of rapid turnover of SD systems. Previous work has demonstrated the Y-linked duplicated copy of *Amh* (*Amhy*) on LG23 as the SD gene of *ONJp* [25]. This has been confirmed by our analyses of Illumina reads generated from male *ONJp* individuals with an XY karyotype. As expected, the XY reads show excessive numbers of SNPs (i.e., differences between the X- and Y-linked alleles or gametologs) along the X chromosome (**Figure 5a**), and also a nearly doubled read coverage of a YY male (derived from crossing the wild-type male with the sexually reversed XY female) indicative of duplication at the region encompassing *Amh* (**Figure 5b**), compared to the surrounding regions, or the patterns derived from female (XX) reads (**Supplementary Fig. S13**). With the same rationale, we used the ZW reads of *OA* and identified LG3 as its sex chromosome pair (**Figure 5c**) with excessive ZW-derived SNPs. We managed to exclude the segregating polymorphic sites and further narrowed down the previously identified SD region (SDR) on the Z chromosome from about 40 Mb long into 0.6Mb, by inspecting the newly produced resequencing data, as well as other published data [22, 48] of different *OA* populations. We first identified the fixed female-specific SNPs or indels that are shared among all the populations which are only concentrated at a 10Mb long region (**Figure 5d**). We then focused on an enclosed region that shows the highest density of female-specific heterozygotes, i.e., the largest differences between the Z and W chromosomes. We genotyped randomly selected candidate sex-linked markers within the region (**Supplementary Table S4**), and found one deletion (**Supplementary Fig. S14**) and one SNP site that are specific to the W chromosomes of all the inspected female *OA* individuals. This candidate SDR spans 620kb and harbors three candidate SD genes, *Banf2, Paics-1*, and *Paics-2*. Intriguingly, these genes show an elevated female vs. male read coverage, suggesting that they are duplicated on the W chromosome (**Figure 5e**).

**Figure 5.**
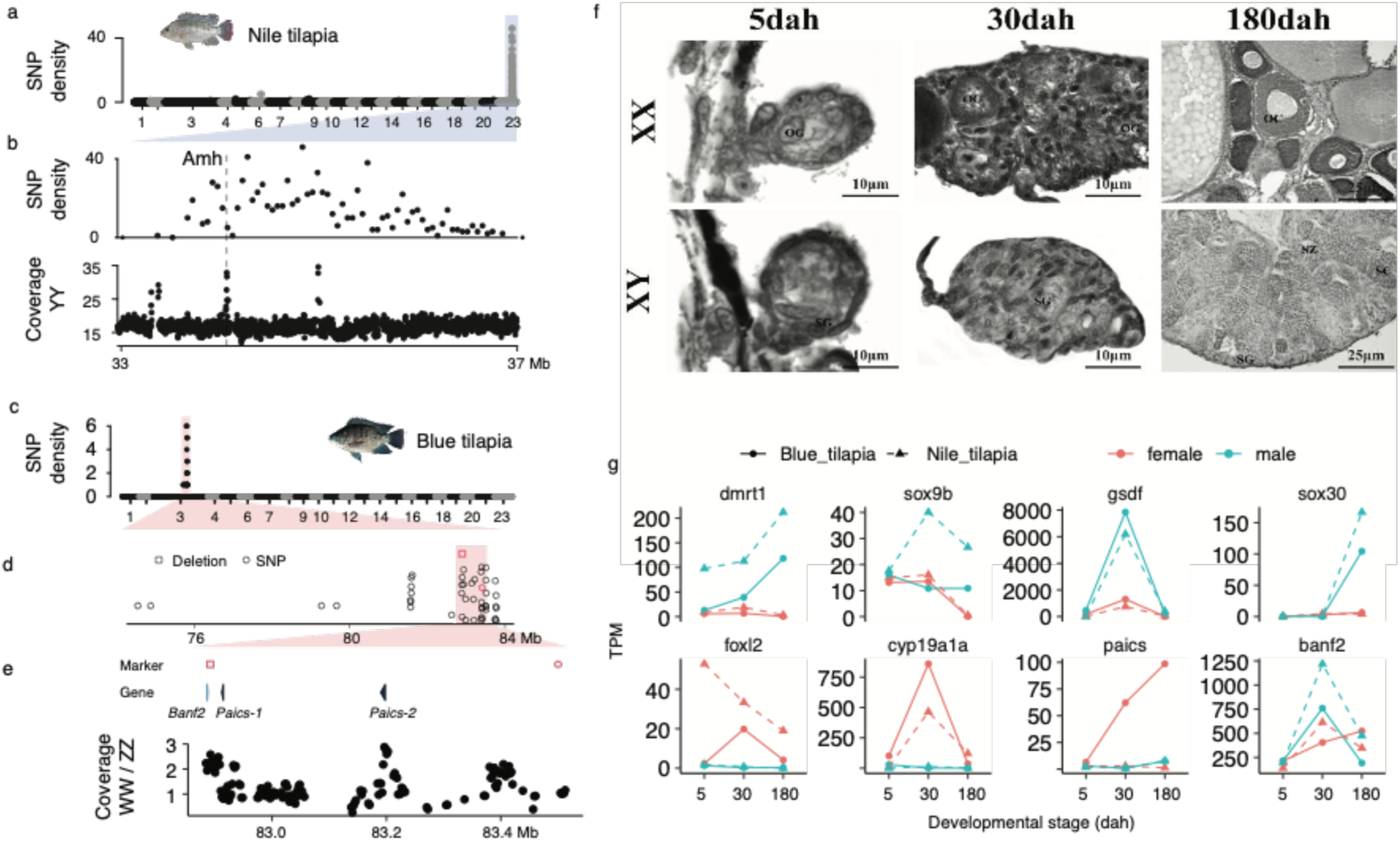
Sex-determining region of Nile tilapia and blue tilapia. a) Distribution of male-specific SNP (number of SNPs per 50k window) in *ONJp*. b) The zoom-in view of the sex-determining region on LG23. The coverage of YY male was calculated for each 5kb window. The location of the sex-determining gene *Amhy* is indicated by a vertical dashed line. c) Distribution of female-specific SNP (number of SNPs per 50kb window) in *OA*. d) The zoom-in view for the sex-determining region showing all female-specific variants. e) The zoom-in view for the region that contains the verified female-specific variants. The verified SNP is highlighted in red. The ratio of coverage of WW female and ZZ male was calculated for every 5kb window. Windows with less than 60% base pairs mapped are not shown. f) We examined oogonia and spermatogonia in the XX and XY gonads of *ONJp* at 5 dah, when no morphological differences can be found between sexes. At 30 dah, oogonia and oocytes can be observed in the XX gonads, indicating the initiation of meiosis. But only spermatogonia can be found in the XY gonad at 30 dah. At 180 dah, the XX gonads display large previtellogenic oocytes, while the XY gonads are characterized by the appearance of spermatogonia, spermatocytes and spermatids. OG, oogonia; SG, spermatogonia; OC, oocytes; SC, spermatocytes; ST, spermatids; SZ, spermatozoa. g) The expression profiles over three stages of gonad development are shown for six known teleost SD genes and two candidate SD genes of blue tilapia.

We hypothesize that a master SD gene, might be expected to show transient sex-specific gene expression during early gonadogenesis, similar to the *Sry* of eutherian mammals[49]. To inspect the candidate SD genes of *OA* for their expression, and also to elucidate the impact of sex chromosome turnovers between *ONJp* and *OA* on their downstream SD pathway genes, we collected the gonad transcriptomes of both sexes from these two species’ corresponding stages. The collected stages span the onset (from 5 days after hatching, or 5-dah), an early (30-dah) and a late (180-dah) stages of gonad differentiation [50], during which the histological differences between gonads of the two sexes become more apparent (**Figure 5f**). Consistently, we found that the numbers of sex-biased genes dramatically increase from 5-dah to the later stages in both sexes of both species (**Supplementary Fig. S15**). In particular, *Paics* have multiple tandem copies at the SDR of both Z (**Figure 4b**) and W (**Figure 5e**) chromosomes of *OA*, and show an increasing ovary-specific expression pattern (**Figure 5g**) through early gonadogenesis in *OA* but not in *ONJp*, suggesting it is likely a candidate female SD gene. The orthologous genes of *Paics* are specifically expressed in gonads of both sexes in *MZ* (**Supplementary Fig. S11**), suggesting that evolution of the potential female-determining function of *Paics* in *OA* might involve suppressing its expression in males. The validation and detailed dissection of *Paics* function require a complete sequence of W chromosomes and more experimental work in future.

The recent transition between the XY chromosomes of *ONJp* and the ZW chromosomes of *OA* is expected to rewire the downstream SD pathways of the two species. To test that, we compared the two species’ gonad expression trajectories of orthologous genes of known vertebrate SD genes: majority of them show a conserved sex-biased temporal expression pattern, but to a different degree, between the two species across the sampled stages (**Supplementary Fig. S16-17**). This suggests that these genes participate in the SD process of both species, but may play a different role because of the turnover of their upstream SD genes. Among them, knockouts in *ONJp* of conserved teleost male-determining genes *Amhy, Gsdf* or *Dmrt1*, or those of female-determining genes *Foxl2* or *Cyp19a1a* all leads to sex reversal [27, 51, 52]. The temporal expression patterns of these genes are consistent with their known hierarchical positions in the SD pathway of *ONJp*: for example, *Dmrt1* has robust male-specific expression and steady upregulation since 5dah, before its validated downstream target genes *Gsdf* [52], *Sox9b* [53] and *Sox30* [54] reaching their peak expression levels specifically in males. This is similar in the female determining pathway between the upstream gene *Foxl2* vs. its downstream target *Cyp19a1a* [27]. Of particular interest is the much higher expression level of *Dmrt1* in *ONJp* than in *OA* in their gonads of 5dah when sex is determined. This is probably because of the origination of new master male SD gene *Amhy* in *ONJp*, whose disruption has been demonstrated to suppress the expression of *Dmrt1* in the male gonads[25]. The increased expression of *Dmrt1* may also account for those of its downstream genes *Sox9b* and *Sox30* in *ONJp* than in *OA*. Interestingly, the upstream female SD gene *Foxl2* is also upregulated in *ONJp* than in *OA*. Since *Dmrt1* and *Foxl2* have a conserved antagonistic relationship during vertebrate SD process that disruption of one would cause the upregulation of the other in the respective sex [55]; an increased expression level of *Foxl2* in *ONJp* female could result from the co-evolution in response to *Dmrt1* in male, or replacement of early female SD role of *Foxl2* in *OA* by the newly evolved candidate SD gene *Paics*.

## Discussion

The great diversity of phenotypes and sex chromosomes generated in a relatively short evolutionary time range makes African cichlids a classic model for studying the mechanisms of species radiation and sex chromosome transitions. Since the release of five representative cichlid genomes over five years ago [3], analyses of more cichlids’ (e.g., those from Lake Malawi [56, 57] and Lake Mweru [58]) and higher qualities of genomes (e.g, that of *ONEg* [22]) have been published, demonstrating the lasting interest in these species. Here we focused on two important aquaculture species from the much less species-rich but much more widely distributed *Oreochromis* genera that have undergone very recent transition between XY and ZW sex chromosome systems. Their high-quality genomes demonstrated by our in-depth analyses of the complex repetitive regions provided novel insights into the genome architecture and sex chromosome evolution of teleosts.

### Chromosome-specific heterochromatin made LG3 a ‘sexy’ chromosome for becoming tilapia sex chromosomes

A few known genes (so-called ‘usual suspects’[59]), for example, *Dmrt1, Amh, Sox3* and *Gsdf* etc. have a conserved role in the vertebrate SD pathway, and frequently evolved to become a master SD gene through point mutations or duplication of one allele in the proto-sex chromosome pair. Their residing ancestral chromosomes or chromosomal fragments (the ‘sexy’ chromosome) are therefore recruited as sex chromosomes more often than other chromosomes [60, 61]. For example, the chicken Z chromosome harboring *Dmrt1* has been independently recruited as sex chromosomes in monotremes and in a gecko species [60]. Substantial proportions of bullfrog sex chromosomes harboring *Sox3* are homologous to the human X chromosome [62]. Among tilapias, LG1 and LG3 are the sexy chromosomes that have been most frequently found as sex chromosomes in all the investigated species [9, 21], although some species have recently evolved SD genes on other LGs (e.g., LG14 of *O. mossambicus* and LG23 of *ONJp*) [25, 45]. In contrast, LG1 and LG3 have not been found as sex chromosomes in other non-tilapia cichlids, except for a sex-linked QTL on LG3 in two Lake Malawi cichlid species[63]. LG23 of *ONJp* became sex chromosomes because of the Y-linked duplication of *Amh* [25]. However, no other ‘usual suspects’ or known master SD genes have been detected within the SDR of LG1 and LG3 [20, 22, 30, 45], suggesting an alternative scenario for recruiting them as sex chromosomes.

In this study, we suggest that tilapia- and chromosome-specific expansion of heterochromatin may have contributed to the more frequent recruitment of LG3 as sex chromosomes. By assembling the nearly complete heterochromatic region and comparison to the Lake Malawi cichlid *MZ*, we inferred that there were two waves of RE (**Figure 3c, Supplementary Fig. S9**). One is probably shared by tilapias and Lake cichlids, and the other is only shared by tilapias on LG3. Both REs dramatically increased the TE content (**Figure 1d**), and decreased the recombination rate across over two thirds of the LG3 region[30], compared to other LGs. The classic model of sex chromosome evolution, as indicated by mammals and birds, hypothesizes that the origination of SD genes would select for suppression of recombination and lead to accumulation of repetitive elements on the Y or W chromosomes[64]. Given the divergence level and the SDR length between sex chromosomes of *OA* are much smaller than those of mammals and birds, it seems more likely a reversed process occurred in which RE and reduction of recombination rate predated and facilitated the emergence of new SD genes on LG3. The abundant repetitive sequences can promote gene duplications or other types of mutations that endow the new SD function to the pre-existing alleles. And the substantial linkage disequilibrium created by the large heterochromatic region probably will further fix the combination of the newly invaded SD locus with other sexually antagonistic loci on the same chromosomes. A similar scenario has been suggested for other cichlids [65, 66] and guppies [67].

### Newcomers of tilapia SD genes

The low sex chromosome divergence level (**Figure 5**) suggested that LG3 as a sex chromosome pair of *OA* evolved very recently, although it requires further confirmation of SDR and candidate SD genes on LG3 of other tilapias (e.g., *O. karongae* and *O. tanganicae*)[10, 45]. We identified the candidate SD genes *Paics* on LG3 of *OA*, which have two Z-linked copies within the SDR. *Paics* genes have an increased copy number on the W chromosome and an ovary-specific expression pattern during gonadogenesis of *OA* but not in *ONJp*, but their human ortholog is ubiquitously expressed across all tissue types without an obvious sex-biased pattern (**Supplementary Fig. S18**). The human *Paics* encodes the phosphoribosylaminoimidazole carboxylase that participates in the purine biosynthesis without sex-related functions. How did *Paics* evolve their ovary-specific expression from the ancestral non-biased expression pattern; and what are the role of the extra W-linked *Paics* copies, if any, during the female SD process of *OA* remain intriguing questions for the future experimental studies. They also require a complete sequence of the W chromosome of *OA*.

Many ‘usual suspects’ or their duplications were identified as the master SD genes in teleosts, e.g., *Amhy* of *ONJp, Dmrt1bY (DMY)* or *Sox3Y* in different medaka species[68-70] etc. Nevertheless, more ‘newcomers’ of master SD genes like *Paics*, i.e., genes that have no previously known SD functions, have now been discovered. For example, the candidate male SD gene of the channel catfish seems to be the male-specific isoform of breast cancer anti-resistance 1 (*BCAR1*) gene [71]. And the male SD gene of rainbow trout *sdY* (*Oncorhynchus mykiss*) is derived from duplication and truncation of an immunity-related gene *irf9 [72]*. It has been recently shown that *sdY* functions by hijacking the female SD regulatory loop between *Foxl2* and *Cyp19a1* to promote testis differentiation[73]. Thus, it is possible that *Paics*, if demonstrated to be a true female SD gene, might similarly interfere with the interaction loop of male SD pathway (e.g., between *Dmrt1* and *Sox9b*) or promote that of the female SD pathway involving *Foxl2* and *Cyp19a1a*. All these key SD genes indeed have a different expression level between *OA* and *ONJp* in the early gonads (**Figure 5g**). Tests of these hypotheses will require transgenic expression of *Paics* genes of *OA* in the females of technically more accessible *ONJp* to evaluate their impact on the known SD genes.

### An important resource for cichlid functional genomic studies and aquaculture

Previous comparison between *ONEg* and *MZ* [30] indicated that the genomic differences between the two species are dominated by intra-rather than inter-chromosomal rearrangements, consistent with a largely conserved karyotype among African cichlids shown by cytogenetic studies [42, 74]. Therefore, the much improved genomes of *ONJp* and *OA* generated by this study in chromosome shape are to provide a high-quality reference for future studies into the patterns and mechanisms of speciation of Lake cichlids. They will be very useful for anchoring the genomes of other cichlid species into chromosomes, and annotating their genes and conserved functional non-coding regulatory elements. More importantly, so far a total of 6 SD genes have been successfully knocked out in *ONJp* [25, 27, 51, 52, 75], with other well-established genetic resources and techniques like antibodies and transgenics, as well as the high-quality genome available now, *ONJp* becomes a promising model for testing the functions of identified genes responsible for the focal phenotypes (e.g., sex determination, body colors) in the future.

The completeness of our new genomes is exemplified by our assembly of complex repetitive regions like the LG3 heterochromatin and rDNA clusters (**Figure 2-3**). The genomic sequences, particularly the fixed sex-specific markers identified here in *ONJp* and *OA* (**Figure 5b,e**) can assist hybridization schemes aiming for producing monosex tilapia fry. For example, these markers can be used to discriminate between sexually reversed ZZ female *OA* vs. the wild-type females, so that the ZZ female can be further crossed with the wild type ZZ male to produce all-male fry, which are preferred over females in aquaculture.

## Conclusion

In this work, we generated and analyzed the chromosome-level genomes of two important aquaculture species *ONJp* and *OA*. We characterized their complex repetitive regions including centromeres, rDNA loci, and the chromosome-specific heterochromatic region on LG3. We showed that the acquisition of LG3 heterochromatin is the result of two episodes of repeat expansions, accompanied by dramatically reduced recombination rate [30] over two thirds of this chromosome. Within the LG3 heterochromatin, we identified a candidate female SD gene in *OA* that showed an ovary-specific expression pattern during the critical stage of sex determination. Overall, our work provides important genomic resources for studying SD mechanisms and genome architectures of tilapias.

## Materials and Methods

### DNA sampling and sequencing

All animal experiments were conducted in accordance with the regulations of the Guide for Care and Use of Laboratory Animals and were approved by the Committee of Laboratory Animal Experimentation at Southwest University. High molecular weight DNAs of *ONJp* (derived from Prof. Nagahama at National Institute for Basic Biology of Japan) and *OA* (from Wuxi Freshwater Fisheries Center in China) were extracted from muscle tissues using a Blood & Cell Culture DNA Midi Kit (Q13343, Qiagen, CA, USA). We also obtained genomic DNAs from *ONJp* with a YY genotype, and *OA* with a WW genotype, by crossing the XY male with the XY female of *ONJp*, and the ZW female with the ZW male of *OA*. The sexually reversed XY female or ZW male individuals were produced by treating fry with the aromatase inhibitor Fadrozole (Novartis)[23] or 17-alpha-ethynylestradiol[76]. We performed the DNA quality and quantity assessment using a Qubit double-stranded DNA HS Assay Kit (Invitrogen, Thermo Fisher Scientific) and an Agilent Bioanalyzer 2100 (Agilent Technologies). For each Nanopore library, approximately 8 µg of gDNAs from the female *ONJp* (XX genotype) and male *OA* (ZZ genotype) were size-selected (10-50 kb) with a Blue Pippin (Sage Science, Beverly, MA), and processed using the Ligation sequencing 1D kit (SQK-LSK108, ONT, UK) according to the manufacturer’s instructions. Libraries were constructed and sequenced on R9.4 FlowCells using the GridION X5 sequencer (ONT, UK) each at the Genome Center of Nextomics (Wuhan, China). To acquire a chromosomal-level assembly of the genome, one gram of gonad tissues collected from the same *ONJp* or *OA* strain of the same genotype was used for Hi-C library construction. The Hi-C experiment consisted of cell crosslinking, cell lysis, chromatin digestion, biotin label, proximity chromatin DNA ligations and DNA purification, which were performed by Annoroad Genomics (Beijing, China) following the standard procedure [77]. The purified and enriched DNA was used for sequencing library construction. Illumina HiSeq X Ten platform (Illumina) was used to perform sequencing with a read length of 150 bp for each end. To identify the candidate SDR, we also performed Illumina sequencing for the YY *ONJp* and WW *OA* individuals with a 250bp library insert size.

### Genome assembly

We used flye (2.3.1) [78] to assemble the Nanopore raw reads, with default parameters. The draft assembly was then polished by Racon (v1.3.1) [79]. To do so, we mapped the raw Nanopore reads using minimap2 (2.15-r905) [80], with options ‘-x map-ont --secondary=no’. We performed Racon polishing for two rounds with default parameters. We then used purge_haplotigs [81] to remove tentative haplotigs (alternative haploid contig). Coverage distribution of Nanopore reads were calculated using the readhist module in purge_haplotigs, after the reads were mapped against the assembly by minimap2 [80]. We used the options ‘-j 80 -s 80’ to decide the classification of haplotigs, and the haplotigs were subsequently removed. The 3D-DNA pipeline (180922) [82] was used to join the contigs into chromosomes. First, we mapped the Hi-C reads against the contigs using Juicer (1.7.6) [83] with default settings. After removing the duplicates, the Hi-C contact map was directly taken as input for 3D-DNA. The parameters were set as ‘--editor-coarse-resolution 500000 --editor-coarse-region 1000000 --editor-saturation-centile 5 -r 0’. We subsequently used Juicebox Assembly Tools [84] to review and manually curate scaffolding errors. We further used Pilon (1.22) [85] to polish the assembly with Illumina sequencing reads. For the ZZ genome (OA), ∼40X sequencing data from a short-insert library was produced for polishing the assembly. Those options were used by Pilon: ‘--minmq 30 --diploid --fix bases,gaps --mindepth 15’. To assess the completeness of the assembled genome, we screened the assembly for BUSCO genes (3.0.2) [86] of actinopterygii. The ‘geno’ model was used with default parameters.

### Genome annotation

We used RepeatModeler (1.0.10) to predict repetitive elements throughout the genome and to classify the repeats based on their similarity to known repeat families. The unclassified repeats were labelled as ‘unknown’. We then combined the newly predicted repeat family with an existing repeat library from RepBase, and used RepeatMasker (4.0.7) to search for repeats in the genomes. We used the MAKER pipeline (2.31.10)] [87] to annotate gene models. The protein sequences of *Oreochromis niloticus* (O_niloticus_UMD_NMBU) [22] and *Maylandia zebra* (M_zebra_UMD2a) [30] were downloaded from NCBI RefSeq as the query to search for homologs. An initial set of gene models were predicted by MAKER with the input of protein sequences alone. We also used Trinity (2.4.0) [88] to assemble the transcriptomes with the parameters ‘--min_glue 5 --path_reinforcement_distance 30 --min_contig_length 300’. We further built a comprehensive transcript database by using the PASA pipeline (2.3.3) [89] with options ‘--stringent_alignment_overlap 30 --ALIGNERS gmap --TRANSDECODER’. Then we fed the MAKER-produced gene models into the PASA pipeline for gene-model polishing (-A --gene_overlap 50).

The known tilapia rDNA sequences were retrieved from NCBI (accession GU289229.1 and MF460358.1) and were mapped against the genome using blastn (2.10.0). For the mapped locus, dotplots were produced by flexidot (1.06) [90] with the parameters -f 1 -k 39 -S 2.

### mRNA sequencing and gene expression analysis

We extracted the total gonad RNAs of each sex of *OA* at 5, 30, and 180 dah (**Supplementary Table S5**) using the Trizol Reagent (Invitrogen, Carlsbad, CA), and eliminated the genomic DNA using DNaseI. RNA qualities were monitored on 1% agarose gels and a Nanodrop spectrophotometer. Illumina sequencing was carried out at Novogene Bioinformatics Technology Co., Ltd., in Beijing, China. Sequencing libraries were constructed using the NEBNext® Ultra™ RNA Library Prep Kit for Illumina® (NEB, USA), according to the manufacturer’s protocol. Index codes were added to attribute sequences to each sample. The prepared libraries were sequenced on an Illumina Hiseq 2500 platform, and 150bp paired-end reads were generated. Gonad transcriptome data of *ONJp* at 5, 30 and 180 dah were from the previous studies [50, 91]. The raw RNA-seq reads were mapped against the genomes using HISAT2 (2.1.0) [92]. The number of reads for each gene was counted using featureCounts (1.6.2) [93] according to the gene model annotation. To quantify the expression levels, read counts were normalised using the TPM (transcripts per million) method. The mean expression levels were calculated for biological replicates.

### Small RNA analysis

The small RNA (sRNA) sequencing data of *ONJp* was retrieved from [94] and [95]. Trimmomatic (0.36) [96] was used to trim adaptors and low-quality bases with the parameter ILLUMINACLIP:adapter:2:30:7 MINLEN:18. The sequencing reads were collapsed using seqcluster (1.2.7) [97] prior to mapping. The aligner bowtie (1.2.1.1) [98] was used to map the reads to the genomes with the parameters --best --strata -k1 - m 1000. The sRNA sequences were compared against small RNA Rfam with cmscan to filter out those that were annotated as microRNA, tRNA, rRNA and other known small RNAs. We further classified the small RNA according to the sequence lengths: piRNA between 26 bp and 32 bp, siRNA between 16 bp and 23 bp. The expression levels of putative piRNA and siRNA loci were quantified by counting read counts at each locus followed by normalization using the TPM method. We used proTRAC (2.4.3) [99] to detect piRNA clusters with default parameters. Prior to running proTRAC, piRNA transcripts were mapped to the genome using the script sRNAmapper.pl following the recommendations by proTRAC.

### Sex determining region

Whole genome resequencing data of multiple males and females (**Supplementary Table S6**) were produced to infer the SDR in both Nile tilapia and blue tilapia. The sequencing reads were mapped against the genome with BWA-mem (0.7.16a). For each sample the variants were called using GATK (3.8.1.0) HaplotypeCaller [100]. The variants were then genotyped together, combining all samples (join calling). The single nucleotide variants were selected and filtered using the criteria QD < 2.0 || FS > 60.0 || MQRankSum < -12.5 || RedPosRankSum < -8.0 || SOR > 3.0 || MQ < 40.0. For *ONJp*, we selected SNPs that were heterozygous (0/1) in males but homozygous (0/0) in females, and for *OA*, we selected female-heterozygous (0/1) but male-homozygous (0/0) SNPs. For pooled resequencing data, we used LoFreq (2.1.2)[101] to call variants, with default parameters. We required the heterozygous SNPs to have an allele frequency between 0.35 and 0.65. The regions contained those sex-linked SNPs were defined as sex-determining regions. For blue tilapia, we further genotyped the SNPs by PCR across the SDR (**Supplementary Table S3**), and discarded the variants that failed to exhibit the sex-linked pattern across all the inspected male and female individuals from different *OA* populations. Sequencing coverage was calculated by Samtools depth (1.3.1) [102] (only for the sites with mapping quality of least 60) followed by calculating the mean coverage in 50k sliding windows along the chromosomes.

### Histological analysis

We sampled XX and XY fish at 5, 30 and 180 dah (days after hatching). Briefly, the fish were anesthetized using an overdose of MS222 (Sigma-Aldrich, St. Louis, USA).

Histological analysis was performed as described [103]. We dissected gonads and fixed the gonads in Bouin’s solution for at least 24 hours at room temperature, dehydrated, and embedded in paraffin. All tissue blocks were sectioned at 5 μm using the Leica microtome (Leica Microsystems, Wetzlar, Germany) and stained with hematoxylin and eosin. Photographs were taken under Olympus BX51 light microscope (Olympus, Tokyo, Japan).

### Data availability

The sequencing reads have been deposited at NCBI SRA, under PRJNA609616. The genome assemblies have been deposited at DDBJ/ENA/GenBank under the accession JAAMTG000000000 and JAAMTF000000000

## Code availability

The codes used in this study have been deposited at https://github.com/lurebgi/tilapiaSexChr.

## Acknowledgement

Q.Z. is supported by the National Natural Science Foundation of China (Grant nos. 31722050 and 31671319), the Natural Science Foundation of Zhejiang Province (LD19C190001), and the European Research Council Starting Grant (Grant agreement 677696). D.W. is supported by the National Natural Science Foundation of China (31861123001 and 31630082), and the National Key Research and Development Program of China (2018YFD0900202). L.X. is supported by uni:docs fellowship from University of Vienna.

## Competing interests

The authors declare that they have no competing interests.

## References

1. Salzburger W, Meyer A: The species flocks of East African cichlid fishes: recent advances in molecular phylogenetics and population genetics. Naturwissenschaften 2004, 91:277–290.

2. Kocher TD: Adaptive evolution and explosive speciation: the cichlid fish model. Nat Rev Genet 2004, 5:288–298.

3. Brawand D, Wagner CE, Li YI, Malinsky M, Keller I, Fan S, Simakov O, Ng AY, Lim ZW, Bezault E, et al: The genomic substrate for adaptive radiation in African cichlid fish. Nature 2014, 513:375–381.

4. Xiao J, Zhong H, Liu Z, Yu F, Luo Y, Gan X, Zhou Y: Transcriptome analysis revealed positive selection of immune-related genes in tilapia. Fish Shellfish Immunol 2015, 44:60–65.

5. Pouyaud L, Agnèse JF: Phylogenetic relationships between 21 species of three tilapiine genera Tilapia, Sarotherodon and Oreochromis using allozyme data. J Fish Biol 1995.

6. Beardmore JA, Mair GC, Lewis RI: Monosex male production in finfish as exemplified by tilapia: applications, problems, and prospects. In Reproductive Biotechnology in Finfish Aquaculture. edited by Lee C-S, Donaldson EM. Amsterdam: Elsevier; 2001: 283–301

7. Mair GC, Abucay JS, Abella TA, Beardmore JA, Skibinski DOF: Genetic manipulation of sex ratio for the large-scale production of all-male tilapia Oreochromis niloticus. Can J Fish Aquat Sci 1997, 54:396–404.

8. Cnaani A, Lee BY, Zilberman N, Ozouf-Costaz C, others: Genetics of sex determination in tilapiine species. Sexualities 2008.

9. Baroiller JF, D’Cotta H, Bezault E, Wessels S, Hoerstgen-Schwark G: Tilapia sex determination: Where temperature and genetics meet. Comp Biochem Physiol A Mol Integr Physiol 2009, 153:30–38.

10. Gammerdinger WJ, Kocher TD: Unusual diversity of sex chromosomes in African cichlid fishes. Genes 2018, 9.

11. Bachtrog D, Mank JE, Peichel CL, Kirkpatrick M, Otto SP, Ashman T-L, Hahn MW, Kitano J, Mayrose I, Ming R, et al: Sex determination: why so many ways of doing it? PLoS Biol 2014, 12:e1001899.

12. Beukeboom LW, Perrin N: The evolution of sex determination. 2014.

13. Mank JE, Avise JC: Evolutionary diversity and turn-over of sex determination in teleost fishes. Sex Dev 2009, 3:60–67.

14. Kottler VA, Schartl M: The Colorful Sex Chromosomes of Teleost Fish. Genes 2018, 9.

15. Kikuchi K, Hamaguchi S: Novel sex-determining genes in fish and sex chromosome evolution. Dev Dyn 2013.

16. Cnaani A: The tilapias’ chromosomes influencing sex determination. Cytogenet Genome Res 2013, 141:195–205.

17. Campos-Ramos R, Harvey SC, Penman DJ: Sex-specific differences in the synaptonemal complex in the genus Oreochromis (Cichlidae). Genetica 2009, 135:325–332.

18. Carrasco LAP, Penman DJ, Bromage N: Evidence for the presence of sex chromosomes in the Nile tilapia (Oreochromis niloticus) from synaptonemal complex analysis of XX, XY and YY genotypes. Aquaculture 1999.

19. Ocalewicz K, Mota-Velasco JC, Campos-Ramos R, others: FISH and DAPI staining of the synaptonemal complex of the Nile tilapia (Oreochromis niloticus) allow orientation of the unpaired region of bivalent 1 observed during …. Chromosome 2009.

20. Gammerdinger WJ, Conte MA, Acquah EA, Roberts RB, Kocher TD: Structure and decay of a proto-Y region in Tilapia, Oreochromis niloticus. BMC Genomics 2014, 15:975.

21. Lee B-Y, Coutanceau J-P, Ozouf-Costaz C, D’Cotta H, Baroiller J-F, Kocher TD: Genetic and physical mapping of sex-linked AFLP markers in Nile tilapia (Oreochromis niloticus). Mar Biotechnol 2011, 13:557–562.

22. Conte MA, Gammerdinger WJ, Bartie KL, Penman DJ, Kocher TD: A high quality assembly of the Nile Tilapia (Oreochromis niloticus) genome reveals the structure of two sex determination regions. BMC Genomics 2017, 18:341.

23. Sun Y-L, Jiang D-N, Zeng S, Hu C-J, Ye K, Yang C, Yang S-J, Li M-H, Wang D-S: Screening and characterization of sex-linked DNA markers and marker-assisted selection in the Nile tilapia (Oreochromis niloticus). Aquaculture 2014, 433:19–27.

24. Eshel O, Shirak A, Dor L, Band M, others: Identification of male-specific amh duplication, sexually differentially expressed genes and microRNAs at early embryonic development of Nile tilapia. Biomed Chromatogr 2014.

25. Li M, Sun Y, Zhao J, Shi H, Zeng S, Ye K, Jiang D, Zhou L, Sun L, Tao W, et al: A tandem duplicate of anti-Müllerian hormone with a missense SNP on the Y chromosome is essential for male sex determination in Nile tilapia, Oreochromis niloticus. PLoS Genet 2015, 11:e1005678.

26. Cáceres G, López ME, Cádiz MI, others: Fine mapping using whole-genome sequencing confirms anti-Müllerian hormone as a major gene for sex determination in farmed Nile tilapia (Oreochromis niloticus L …. G3: Genes, Genomes 2019.

27. Li M-H, Yang H-H, Li M-R, Sun Y-L, Jiang X-L, Xie Q-P, Wang T-R, Shi H-J, Sun L-N, Zhou L-Y, Wang D-S: Antagonistic roles of Dmrt1 and Foxl2 in sex differentiation via estrogen production in tilapia as demonstrated by TALENs. Endocrinology 2013, 154:4814–4825.

28. Wang D-S, Kobayashi T, Zhou L-Y, Paul-Prasanth B, Ijiri S, Sakai F, Okubo K, Morohashi K-i, Nagahama Y: Foxl2 up-regulates aromatase gene transcription in a female-specific manner by binding to the promoter as well as interacting with Ad4 binding protein/steroidogenic factor 1. Molecular Endocrinology 2007, 21:712–725.

29. Conte MA, Kocher TD: An improved genome reference for the African cichlid, Metriaclima zebra. BMC Genomics 2015, 16:724.

30. Conte MA, Joshi R, Moore EC, Nandamuri SP, Gammerdinger WJ, Roberts RB, Carleton KL, Lien S, Kocher TD: Chromosome-scale assemblies reveal the structural evolution of African cichlid genomes. Gigascience 2019, 8.

31. Foresti F, Oliveira C, Galetti Junior PM, Almeida-Toledo LF: Synaptonemal complex analysis in spermatocytes of tilapia, Oreochromis niloticus (Pisces, Cichlidae). Genome 1993, 36:1124–1128.

32. Nikaido M, Suzuki H, Toyoda A, Fujiyama A, Hagino-Yamagishi K, Kocher TD, Carleton K, Okada N: Lineage-specific expansion of vomeronasal type 2 receptor-like (OlfC) genes in cichlids may contribute to diversification of amino acid detection systems. Genome Biol Evol 2013, 5:711–722.

33. Martins C, Oliveira C, Wasko AP, Wright JM: Physical mapping of the Nile tilapia (Oreochromis niloticus) genome by fluorescent in situ hybridization of repetitive DNAs to metaphase chromosomes—a review. Aquaculture 2004, 231:37–49.

34. Ferreira IA, Poletto AB, Kocher TD, Mota-Velasco JC, Penman DJ, Martins C: Chromosome evolution in African cichlid fish: contributions from the physical mapping of repeated DNAs. Cytogenet Genome Res 2010, 129:314–322.

35. Chew JSK, Oliveira C, Wright JM, Dobson MJ: Molecular and cytogenetic analysis of the telomeric (TTAGGG)n repetitive sequences in the Nile tilapia, Oreochromis niloticus (Teleostei: Cichlidae). Chromosoma 2002, 111:45–52.

36. Franck JP, Wright JM, McAndrew BJ: Genetic variability in a family of satellite DNAs from tilapia (Pisces: Cichlidae). Genome 1992, 35:719–725.

37. Franck JP, Kornfield I, Wright JM: The utility of SATA satellite DNA sequences for inferring phylogenetic relationships among the three major genera of tilapiine cichlid fishes. Mol Phylogenet Evol 1994, 3:10–16.

38. Oliveira C, Wright JM: Molecular cytogenetic analysis of heterochromatin in the chromosomes of tilapia, Oreochromis niloticus (Teleostei: Cichlidae). Chromosome Res 1998, 6:205–211.

39. Muller H, Gil J, Drinnenberg IA: The impact of centromeres on spatial genome architecture. Trends in Genetics 2019, 35:565–578.

40. Ichikawa K, Tomioka S, Suzuki Y, Nakamura R, Doi K, Yoshimura J, Kumagai M, Inoue Y, Uchida Y, Irie N, et al: Centromere evolution and CpG methylation during vertebrate speciation. Nat Commun 2017, 8:1833.

41. Supiwong W, Tanomtong A, Supanuam P, Seetapan K, Khakhong S, Sanoamuang L-O: Chromosomal characteristic of Nile tilapia (Oreochromis niloticus) from mitotic and meiotic cell division by T-Lymphocyte cell culture. CYTOLOGIA 2013, 78:9–14.

42. Poletto AB, Ferreira IA, Cabral-de-Mello DC, Nakajima RT, Mazzuchelli J, Ribeiro HB, Venere PC, Nirchio M, Kocher TD, Martins C: Chromosome differentiation patterns during cichlid fish evolution. BMC Genet 2012, 13:2.

43. Symonová R: Integrative rDNAomics—importance of the oldest repetitive fraction of the eukaryote genome. Genes 2019, 10:345.

44. Willard HF, Waye JS: Hierarchical order in chromosome-specific human alpha satellite DNA. Trends Genet 1987, 3:192–198.

45. Gammerdinger WJ, Conte MA, Sandkam BA, others: Characterization of sex chromosomes in three deeply diverged species of Pseudocrenilabrinae (Teleostei: Cichlidae). Hydrobiologia 2019.

46. Bolívar P, Mugal CF, Nater A, Ellegren H: Recombination rate variation modulates gene sequence evolution mainly via GC-biased gene conversion, not Hill-Robertson interference, in an avian system. Mol Biol Evol 2016, 33:216–227.

47. Senti K-A, Brennecke J: The piRNA pathway: a fly’s perspective on the guardian of the genome. Trends Genet 2010, 26:499–509.

48. Shirak A, Zak T, Dor L, Benet-Perlberg A, Weller JI, Ron M, Seroussi E: Quantitative trait loci on LGs 9 and 14 affect the reproductive interaction between two Oreochromis species, O. niloticus and O. aureus. Heredity 2019, 122:341–353.

49. Kashimada K, Koopman P: Sry: the master switch in mammalian sex determination. Development 2010, 137:3921–3930.

50. Tao W, Yuan J, Zhou L, Sun L, Sun Y, Yang S, Li M, Zeng S, Huang B, Wang D: Characterization of gonadal transcriptomes from Nile tilapia (Oreochromis niloticus) reveals differentially expressed genes. PLoS One 2013, 8:e63604.

51. Zhang X, Li M, Ma H, Liu X, Shi H, Li M, others: Mutation of foxl2 or cyp19a1a results in female to male sex reversal in XX Nile tilapia. Endocrinology 2017.

52. Jiang D-N, Yang H-H, Li M-H, Shi H-J, Zhang X-B, Wang D-S: gsdf is a downstream gene of dmrt1 that functions in the male sex determination pathway of the Nile tilapia. Mol Reprod Dev 2016, 83:497–508.

53. Wei L, Li X, Li M, Tang Y, Wei J, Wang D: Dmrt1 directly regulates the transcription of the testis-biased Sox9b gene in Nile tilapia (Oreochromis niloticus). Gene 2019, 687:109–115.

54. Tang Y, Li X, Xiao H, Li M, Li Y, Wang D, Wei L: Transcription of the Sox30 Gene Is Positively Regulated by Dmrt1 in Nile Tilapia. International Journal of Molecular Sciences 2019, 20:5487.

55. Lin Y-T, Capel B: Cell fate commitment during mammalian sex determination. Current Opinion in Genetics & Development 2015, 32:144–152.

56. Malinsky M, Svardal H, Tyers AM, Miska EA, Genner MJ, Turner GF, Durbin R: Whole-genome sequences of Malawi cichlids reveal multiple radiations interconnected by gene flow. Nat Ecol Evol 2018, 2:1940–1955.

57. Svardal H, Quah FX, Malinsky M, Ngatunga BP, Miska EA, Salzburger W, Genner MJ, Turner GF, Durbin R: Ancestral hybridization facilitated species diversification in the Lake Malawi cichlid fish adaptive radiation. Molecular Biology and Evolution 2019.

58. Meier JI, Stelkens RB, Joyce DA, Mwaiko S, Phiri N, Schliewen UK, Selz OM, Wagner CE, Katongo C, Seehausen O: The coincidence of ecological opportunity with hybridization explains rapid adaptive radiation in Lake Mweru cichlid fishes. Nat Commun 2019, 10:5391.

59. Herpin A, Schartl M: Plasticity of gene-regulatory networks controlling sex determination: of masters, slaves, usual suspects, newcomers, and usurpators. EMBO Rep 2015, 16:1260–1274.

60. O’Meally D, Ezaz T, Georges A, Sarre SD, Graves JAM: Are some chromosomes particularly good at sex? Insights from amniotes. Chromosome Res 2012, 20:7–19.

61. Graves JAM, Marshall Graves JA, Peichel CL: Are homologies in vertebrate sex determination due to shared ancestry or to limited options? Genome Biology 2010, 11:205.

62. Denton RD, Kudra RS, Malcom JW, Du Preez L, Malone JH: The African Bullfrog (Pyxicephalus adspersus) genome unites the two ancestral ingredients for making vertebrate sex chromosomes.

63. Parnell NF, Streelman JT: Genetic interactions controlling sex and color establish the potential for sexual conflict in Lake Malawi cichlid fishes. Heredity 2013, 110:239–246.

64. Charlesworth D, Charlesworth B, Marais G: Steps in the evolution of heteromorphic sex chromosomes. Heredity 2005, 95:118–128.

65. Roberts RB, Ser JR, Kocher TD: Sexual Conflict Resolved by Invasion of a Novel Sex Determiner in Lake Malawi Cichlid Fishes. Science 2009, 326:998–1001.

66. Ser JR, Roberts RB, Kocher TD: Multiple interacting loci control sex determination in lake Malawi cichlid fish. Evolution 2010, 64:486–501.

67. Bergero R, Gardner J, Bader B, Yong L, Charlesworth D: Exaggerated heterochiasmy in a fish with sex-linked male coloration polymorphisms. Proceedings of the National Academy of Sciences 2019, 116:6924–6931.

68. Nanda I, Kondo M, Hornung U, Asakawa S, Winkler C, Shimizu A, Shan Z, Haaf T, Shimizu N, Shima A, et al: A duplicated copy of DMRT1 in the sex-determining region of the Y chromosome of the medaka, Oryzias latipes. Proceedings of the National Academy of Sciences 2002, 99:11778–11783.

69. Matsuda M, Nagahama Y, Shinomiya A, Sato T, Matsuda C, Kobayashi T, Morrey CE, Shibata N, Asakawa S, Shimizu N, et al: DMY is a Y-specific DM-domain gene required for male development in the medaka fish. Nature 2002, 417:559–563.

70. Takehana Y, Matsuda M, Myosho T, Suster ML, Kawakami K, Shin-I T, Kohara Y, Kuroki Y, Toyoda A, Fujiyama A, et al: Co-option of Sox3 as the male-determining factor on the Y chromosome in the fish Oryzias dancena. Nature Communications 2014, 5.

71. Bao L, Tian C, Liu S, Zhang Y, Elaswad A, Yuan Z, Khalil K, Sun F, Yang Y, Zhou T, et al: The Y chromosome sequence of the channel catfish suggests novel sex determination mechanisms in teleost fish. BMC Biology 2019, 17.

72. Yano A, Guyomard R, Nicol B, Jouanno E, Quillet E, Klopp C, Cabau C, Bouchez O, Fostier A, Guiguen Y: An immune-related gene evolved into the master sex-determining gene in rainbow trout, Oncorhynchus mykiss. Curr Biol 2012, 22:1423–1428.

73. Bertho S, Herpin A, Branthonne A, Jouanno E, Yano A, Nicol B, Muller T, Pannetier M, Pailhoux E, Miwa M, et al: The unusual rainbow trout sex determination gene hijacked the canonical vertebrate gonadal differentiation pathway. Proc Natl Acad Sci U S A 2018, 115:12781–12786.

74. Mazzuchelli J, Kocher TD, Yang F, Martins C: Integrating cytogenetics and genomics in comparative evolutionary studies of cichlid fish. BMC Genomics 2012, 13:463.

75. Xie Q-P, He X, Sui Y-N, Chen L-L, Sun L-N, Wang D-S: Haploinsufficiency of SF-1 Causes Female to Male Sex Reversal in Nile Tilapia, Oreochromis niloticus. Endocrinology 2016, 157:2500–2514.

76. Hopkins KD, Shelton WL, Engle CR: Estrogen sex-reversal of Tilapia aurea. Aquaculture 1979, 18:263–268.

77. Lieberman-Aiden E, van Berkum NL, Williams L, Imakaev M, Ragoczy T, Telling A, Amit I, Lajoie BR, Sabo PJ, Dorschner MO, et al: Comprehensive mapping of long-range interactions reveals folding principles of the human genome. Science 2009, 326:289–293.

78. Kolmogorov M, Yuan J, Lin Y, Pevzner PA: Assembly of long, error-prone reads using repeat graphs. Nat Biotechnol 2019, 37:540–546.

79. Vaser R, Sović I, Nagarajan N, Šikić M: Fast and accurate de novo genome assembly from long uncorrected reads. Genome Res 2017, 27:737–746.

80. Li H: Minimap2: pairwise alignment for nucleotide sequences. Bioinformatics 2018, 34:3094–3100.

81. Roach MJ, Schmidt SA, Borneman AR: Purge Haplotigs: allelic contig reassignment for third-gen diploid genome assemblies. BMC Bioinformatics 2018, 19:460.

82. Dudchenko O, Batra SS, Omer AD, Nyquist SK, Hoeger M, Durand NC, Shamim MS, Machol I, Lander ES, Aiden AP, Aiden EL: De novo assembly of the Aedes aegypti genome using Hi-C yields chromosome-length scaffolds. Science 2017, 356:92–95.

83. Durand NC, Shamim MS, Machol I, Rao SSP, Huntley MH, Lander ES, Aiden EL: Juicer provides a one-click system for analyzing loop-resolution Hi-C experiments. Cell Syst 2016, 3:95–98.

84. Dudchenko O, Shamim MS, Batra SS, Durand NC, Musial NT, Mostofa R, Pham M, St Hilaire BG, Yao W, Stamenova E, et al: The Juicebox Assembly Tools module facilitates de novo assembly of mammalian genomes with chromosome-length scaffolds for under $1000.

85. Walker BJ, Abeel T, Shea T, Priest M, Abouelliel A, Sakthikumar S, Cuomo CA, Zeng Q, Wortman J, Young SK, Earl AM: Pilon: an integrated tool for comprehensive microbial variant detection and genome assembly improvement. PLoS One 2014, 9:e112963.

86. Seppey M, Manni M, Zdobnov EM: BUSCO: Assessing Genome Assembly and Annotation Completeness. Methods Mol Biol 2019, 1962:227–245.

87. Cantarel BL, Korf I, Robb SMC, Parra G, Ross E, Moore B, Holt C, Sánchez Alvarado A, Yandell M: MAKER: an easy-to-use annotation pipeline designed for emerging model organism genomes. Genome Res 2008, 18:188–196.

88. Grabherr MG, Haas BJ, Yassour M, Levin JZ, Thompson DA, Amit I, Adiconis X, Fan L, Raychowdhury R, Zeng Q, et al: Full-length transcriptome assembly from RNA-Seq data without a reference genome. Nature Biotechnology 2011, 29:644–652.

89. Haas BJ, Delcher AL, Mount SM, Wortman JR, Smith RK Jr., Hannick LI, Maiti R, Ronning CM, Rusch DB, Town CD, et al: Improving the Arabidopsis genome annotation using maximal transcript alignment assemblies. Nucleic Acids Res 2003, 31:5654–5666.

90. Seibt KM, Schmidt T, Heitkam T: FlexiDot: highly customizable, ambiguity-aware dotplots for visual sequence analyses. Bioinformatics 2018, 34:3575–3577.

91. Tao W, Chen J, Tan D, Yang J, Sun L, Wei J, Conte MA, Kocher TD, Wang D: Transcriptome display during tilapia sex determination and differentiation as revealed by RNA-Seq analysis. BMC Genomics 2018, 19:363.

92. Kim D, Paggi JM, Park C, Bennett C, Salzberg SL: Graph-based genome alignment and genotyping with HISAT2 and HISAT-genotype. Nature Biotechnology 2019, 37:907–915.

93. Liao Y, Smyth GK, Shi W: featureCounts: an efficient general purpose program for assigning sequence reads to genomic features. Bioinformatics 2014, 30:923–930.

94. Qiang J, Bao WJ, Tao FY, He J, Li XH, Xu P, Sun LY: The expression profiles of miRNA-mRNA of early response in genetically improved farmed tilapia (Oreochromis niloticus) liver by acute heat stress. Sci Rep 2017, 7:8705.

95. Tao W, Sun L, Shi H, Cheng Y, Jiang D, Fu B, Conte MA, Gammerdinger WJ, Kocher TD, Wang D: Integrated analysis of miRNA and mRNA expression profiles in tilapia gonads at an early stage of sex differentiation. BMC Genomics 2016, 17:328.

96. Bolger AM, Lohse M, Usadel B: Trimmomatic: a flexible trimmer for Illumina sequence data. Bioinformatics 2014, 30:2114–2120.

97. Pantano L, Estivill X, Martí E: A non-biased framework for the annotation and classification of the non-miRNA small RNA transcriptome. Bioinformatics 2011, 27:3202–3203.

98. Langmead B, Trapnell C, Pop M, Salzberg SL: Ultrafast and memory-efficient alignment of short DNA sequences to the human genome. Genome Biol 2009, 10:R25.

99. Rosenkranz D, Zischler H: proTRAC - a software for probabilistic piRNA cluster detection, visualization and analysis. BMC Bioinformatics 2012, 13:1–10.

100. Poplin R, Ruano-Rubio V, DePristo MA, Fennell TJ, Carneiro MO, Van der Auwera GA, Kling DE, Gauthier LD, Levy-Moonshine A, Roazen D, et al: Scaling accurate genetic variant discovery to tens of thousands of samples.

101. Wilm A, Aw PP, Bertrand D, Yeo GH, Ong SH, Wong CH, Khor CC, Petric R, Hibberd ML, Nagarajan N: LoFreq: a sequence-quality aware, ultra-sensitive variant caller for uncovering cell-population heterogeneity from high-throughput sequencing datasets. Nucleic Acids Res 2012, 40:11189–11201.

102. Li H, Handsaker B, Wysoker A, Fennell T, Ruan J, Homer N, Marth G, Abecasis G, Durbin R, Genome Project Data Processing S: The Sequence Alignment/Map format and SAMtools. Bioinformatics 2009, 25:2078–2079.

103. Yan L, Feng H, Wang F, Lu B, Liu X, Sun L, Wang D: Establishment of three estrogen receptors (esr1, esr2a, esr2b) knockout lines for functional study in Nile tilapia. J Steroid Biochem Mol Biol 2019, 191:105379.

