## Supplementary Figures for "High-quality chromosome-level genomes of two tilapia species reveal their evolution of repeat sequences and sex chromosomes"

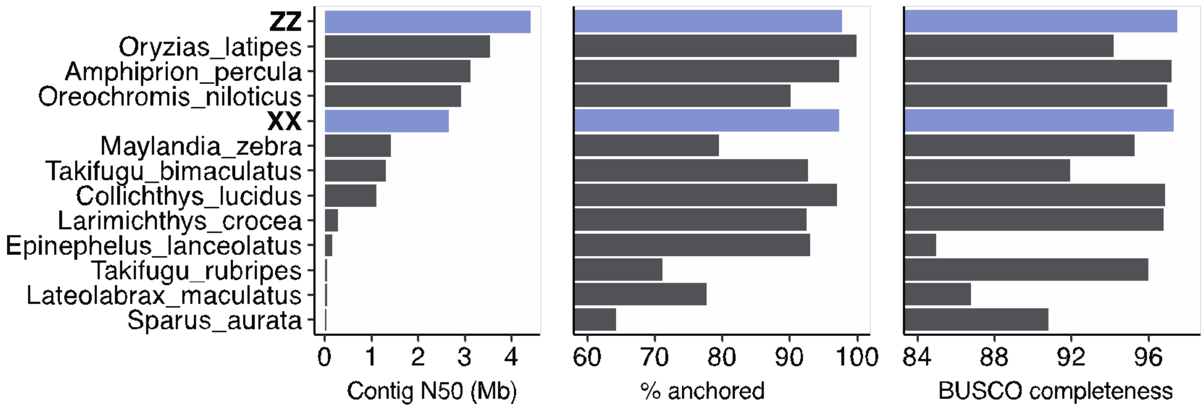


**Supplementary Figure S1 Statistics of genome assemblies at the chromosome scale of teleosts.** Contig N50, the percentage of sequences anchored into chromosome, and genome completeness (BUSCO) were compared among *OA* (ZZ), *ONJp* (XX) and other chromosomal assemblies of teleost genomes.

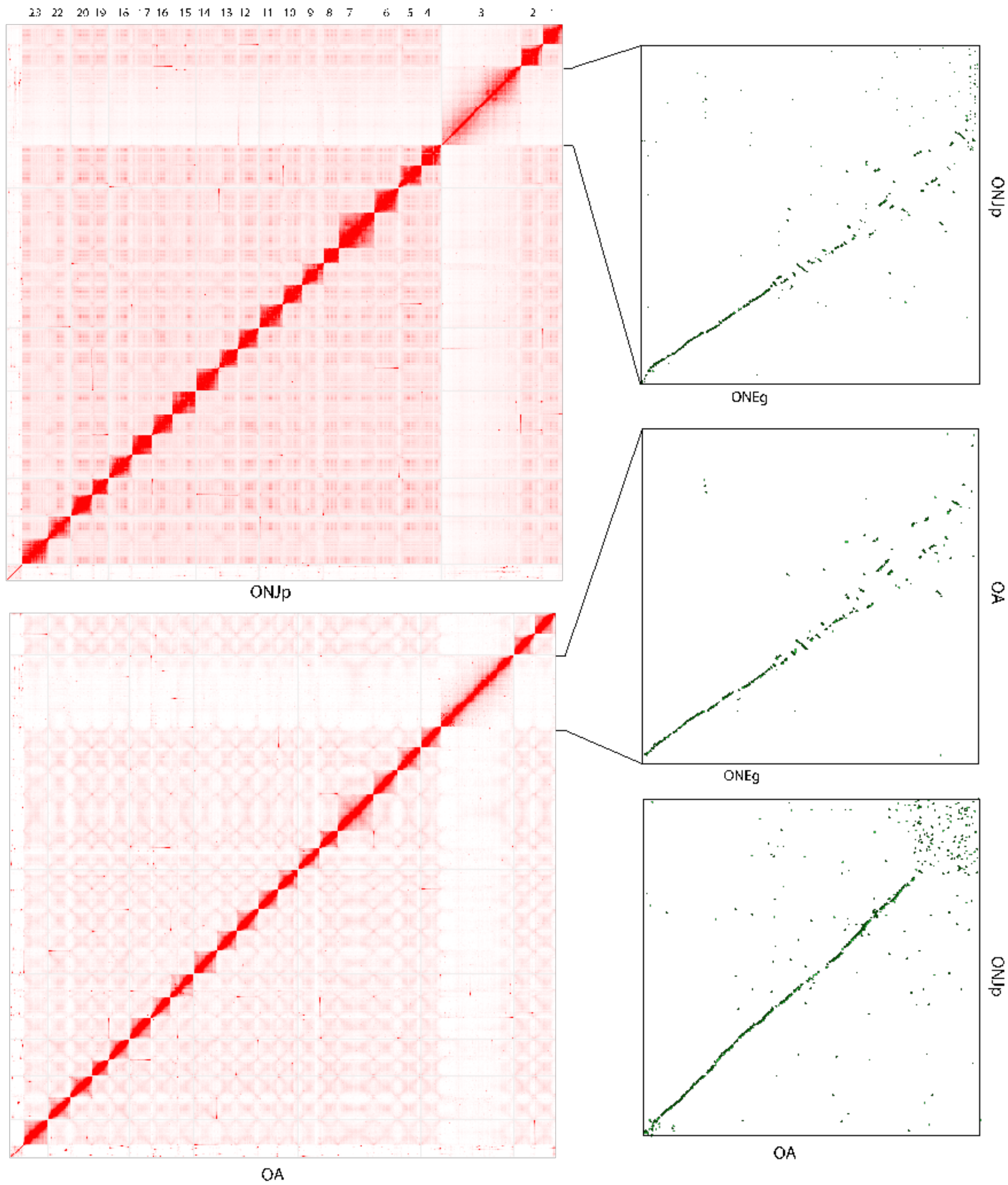


**Supplementary Figure S2 Chromosomal assemblies of tilapia species** The Hi-C contact map produced by Juicebox are shown for both tilapia genomes assembled in this study. The LG3 of each genome is the largest chromosome, and the assembled size is much larger compared with a previous Nile tilapia genome (*ONEg*), demonstrated by the dotplot of synteny (in the right panels)

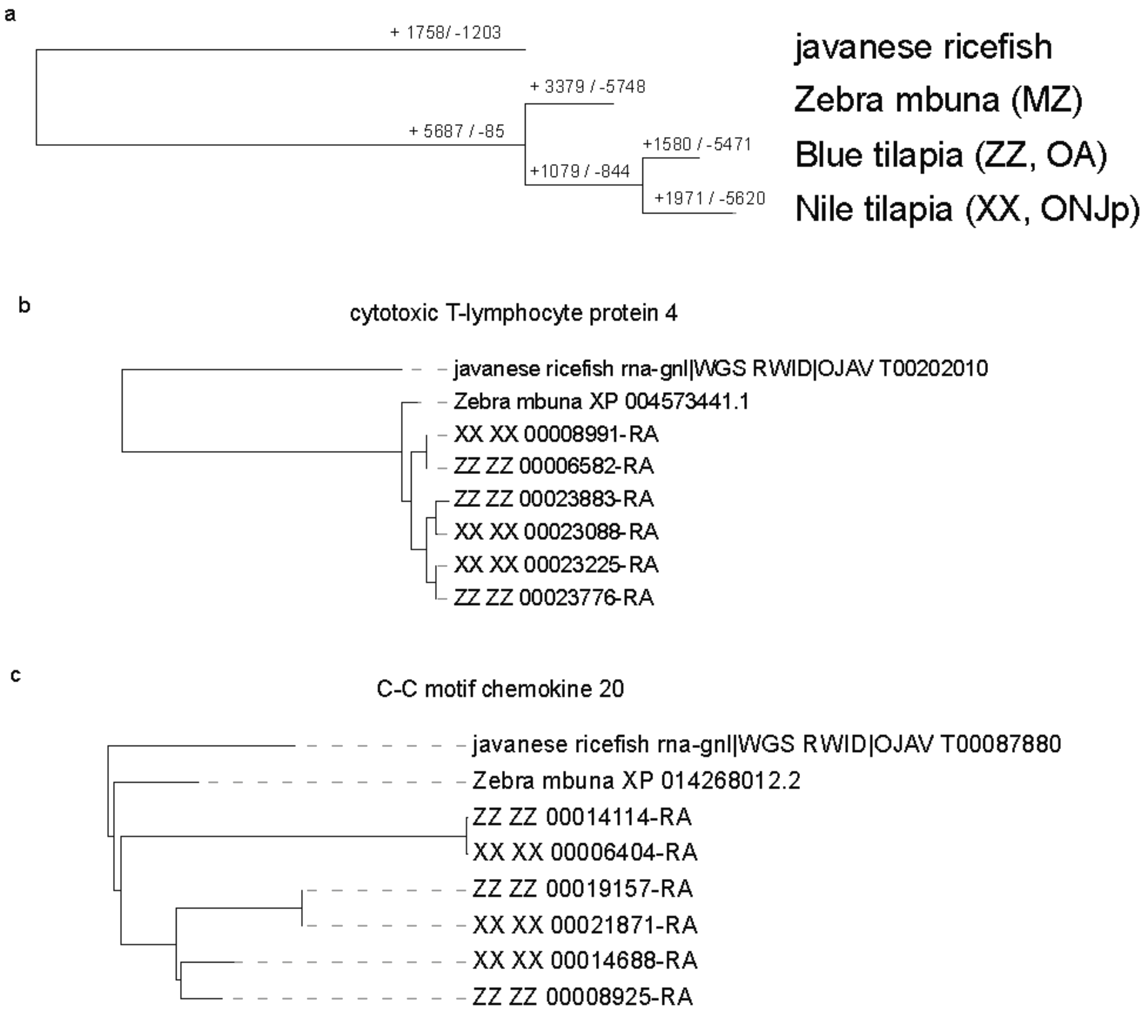


**Supplementary Figure S3 Gene family evolution of tilapias.** a) The orthologous groups were constructed by orthofinder and gene loss and gain events at each phylogenetic node were reconstructed by Notung (2.9.1). b-c) Tilapia-specific duplication of two immune-related genes were shown for demonstration. The reconstructed phylogenetic trees were retrieved from orthofinder outputs.

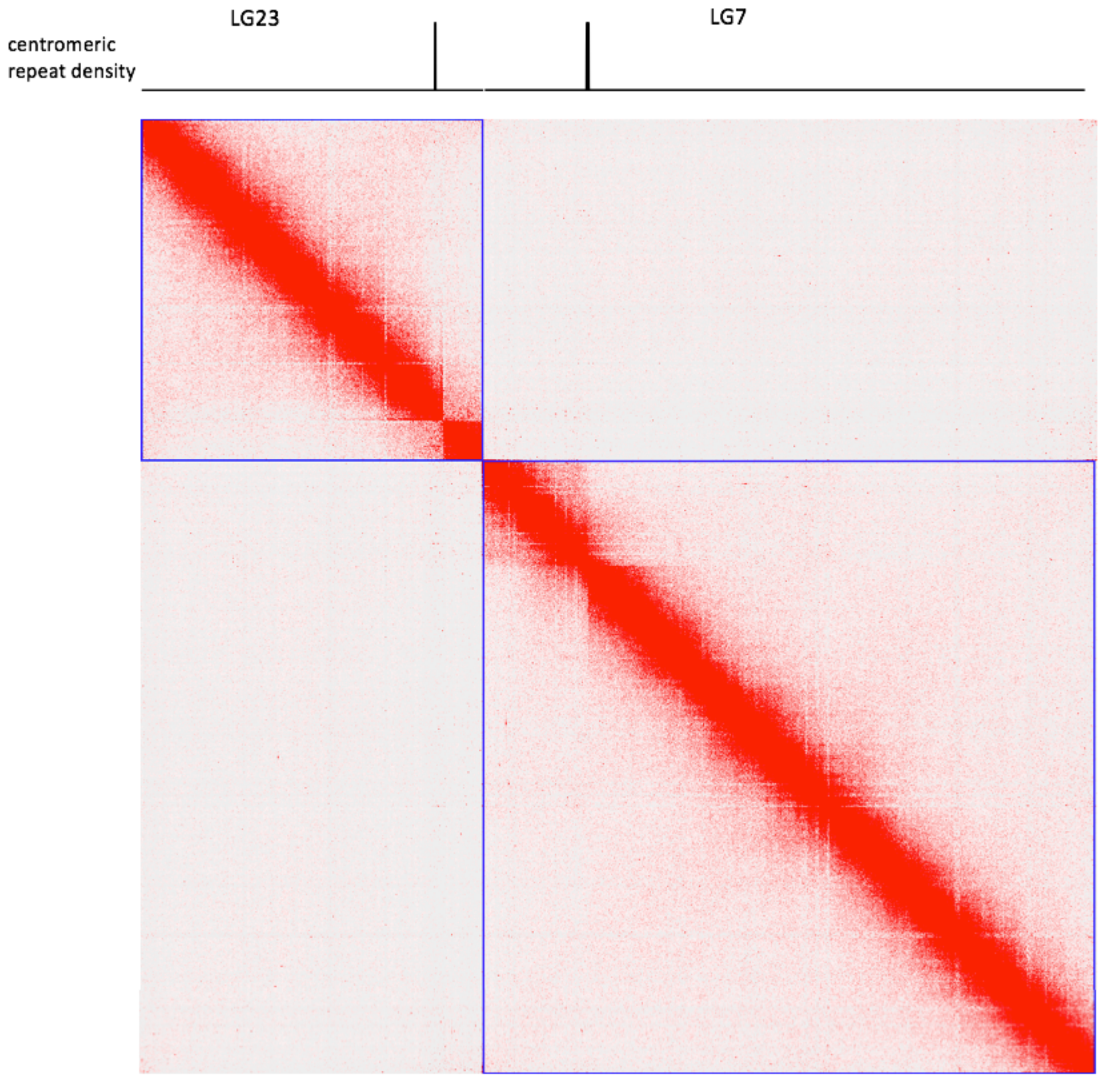


**Supplementary Figure S4 Colocalization of putative centromeres with the boundary between intrachromosomal interaction domains.** The locations of putative centromeres on LG23 and LG7 (those two chromosomes were used for demonstration) are indicated by the peaks of centromeric repeats (SATA monomers) in the upper panel. The Hi-C contact map (produced by Juicebox) of those two chromosomes is shown in the lower panel. The blue lines represent chromosome boundaries.

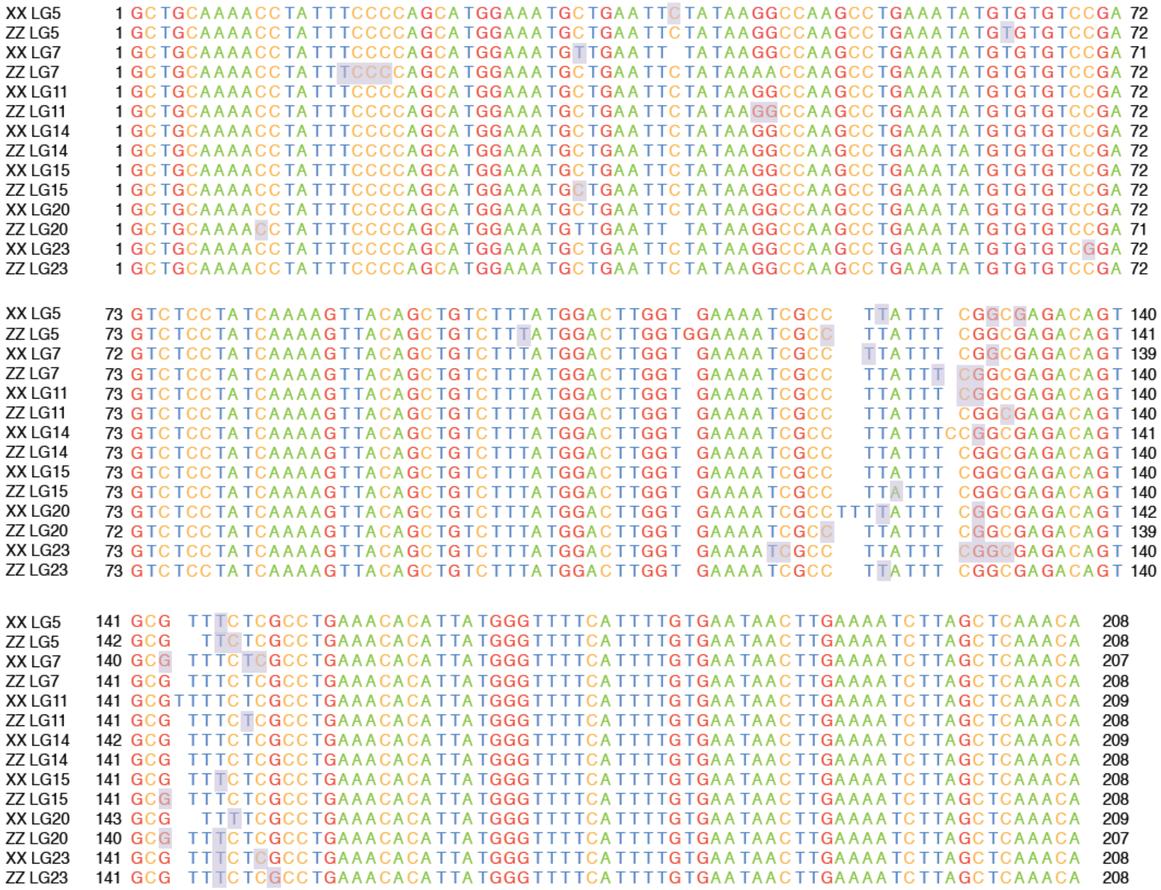


**Supplementary Figure S5 Alignments of SATA monomers of different genomic locations.** The alignments of consensus centromeric monomers from seven chromosomes of Nile tilapia and blue tilapia. The sites in grey background indicate intrachromosomal polymorphisms which also often show interchromosomal variations.

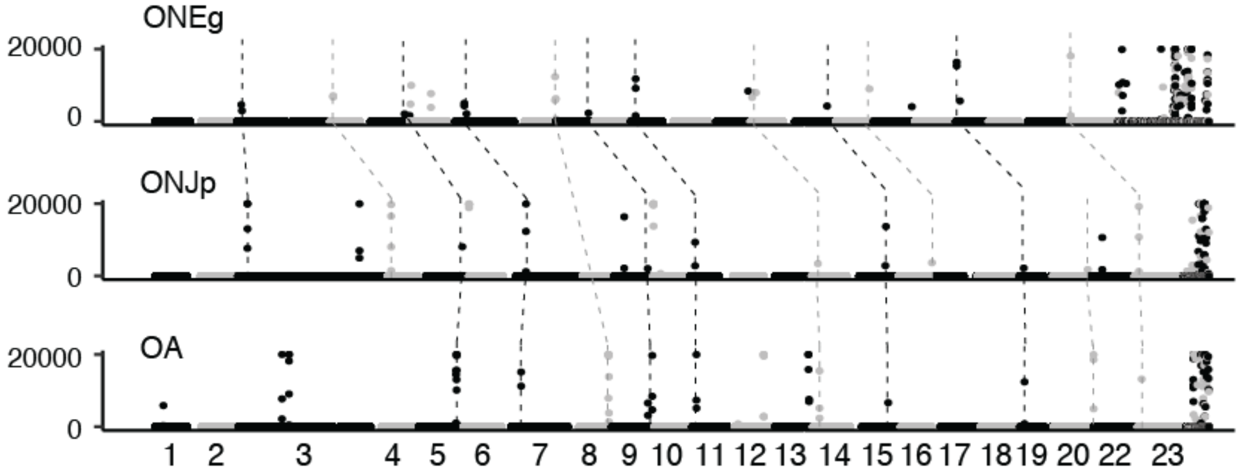


**Supplementary Figure S6 Alignment of putative centromeric positions between tilapia species.** Each dot represents the density of centromeric repeats over 50k windows along the chromosomes. The centromeres that align across species are connected by dashed lines.

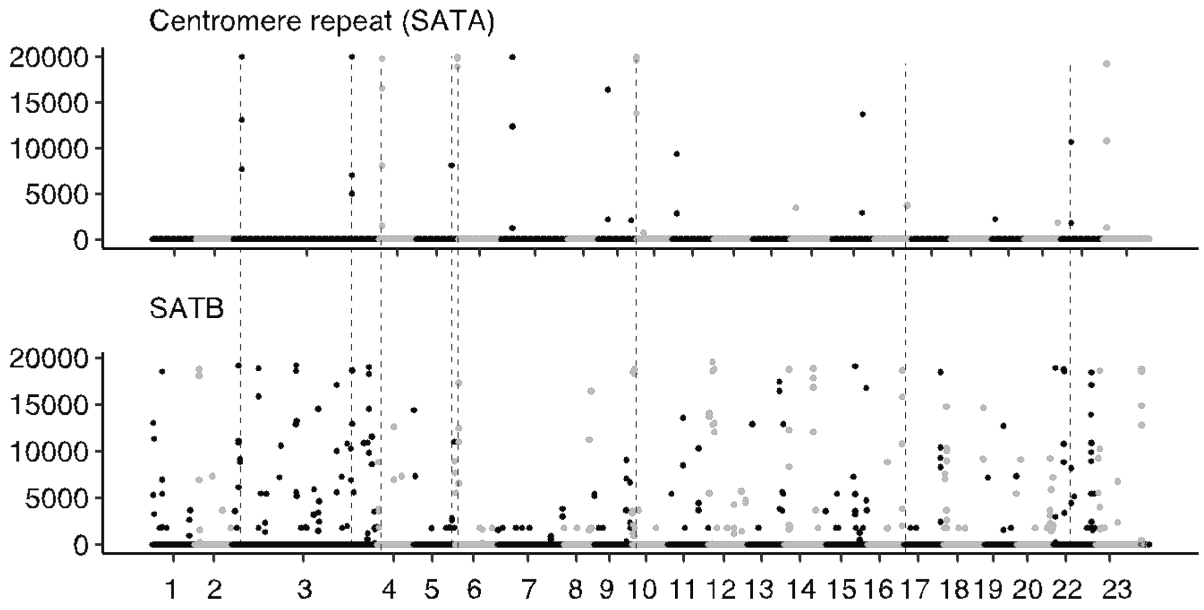


**Supplementary Figure S7 Colocalization of SATA and STAB satellite sequences in *ONJp*.** Each dot represents the density of satellite repeats over 50k windows along the chromosomes. When SATA and SATB repeats co-localize, they are connected with a dashed line.

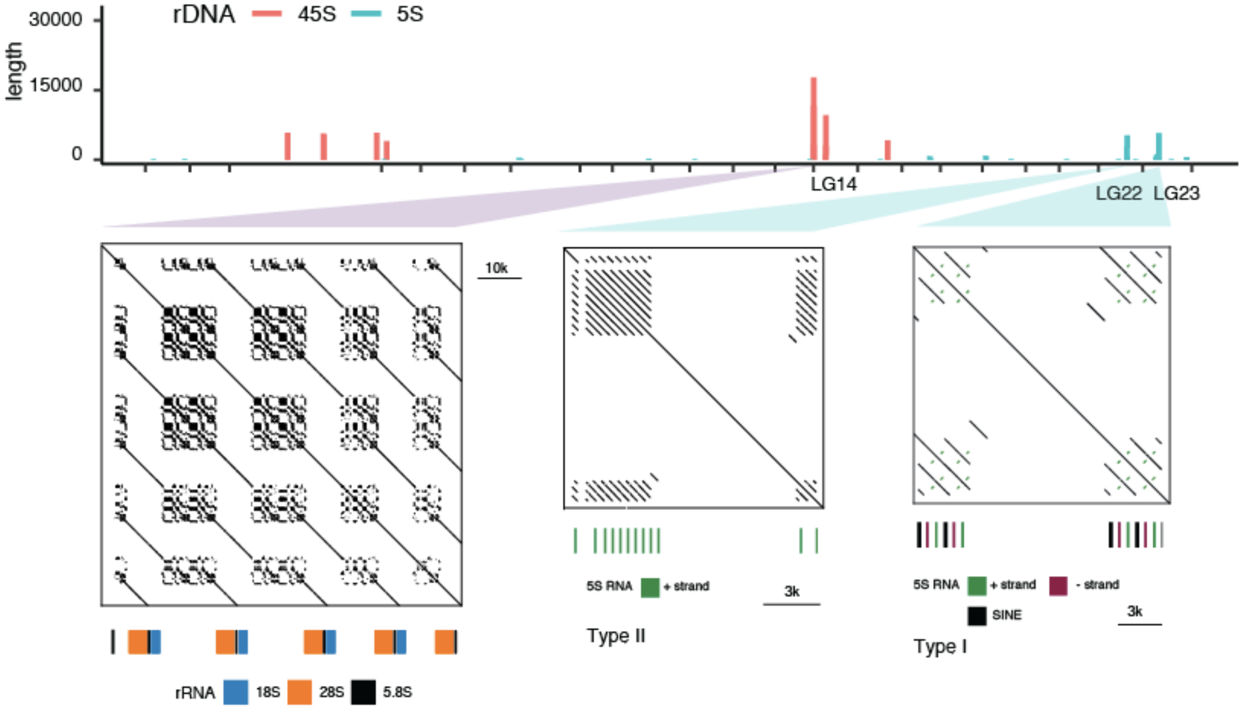


**Supplementary Figure S8 rDNA loci of Nile tilapia (*ONJp*).** The length of rDNA sequence per 50k along the chromosomes of Nile tilapia in the upper panel. One 45S locus on LG14 that contains an array of 4 45S gene loci is demonstrated in a zoom-in view. The 5S locus on LG22 contains a 5S array of type II rRNA and the 5S locus on LG23 contains a 5S array of type I rRNA.

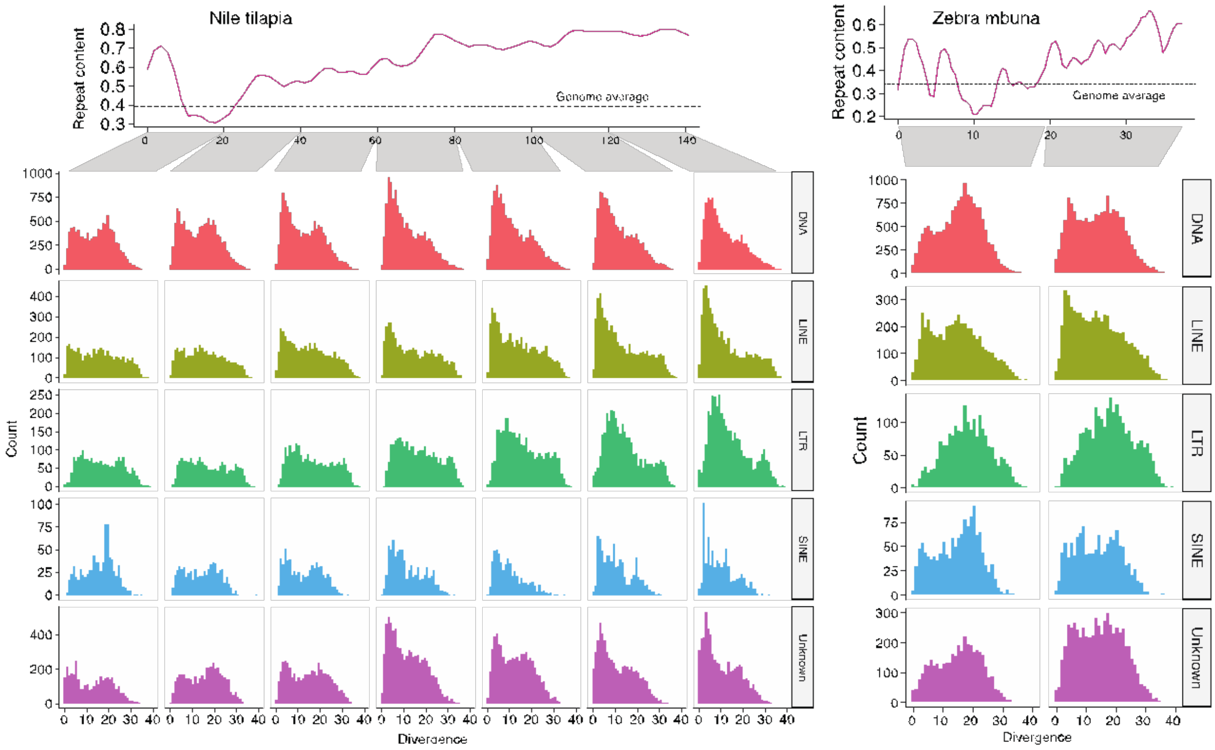


**Supplementary Figure S9 The distributions of sequence divergence of repeats along the LG3 in *ONJp* and *MZ*.** The LG3 was divided into several 20M windows, each window showing the distribution of sequence divergence of different classes of repeats. The 5' end part of LG3 in ONJp tends to have a larger portion of young repeats with lower divergence.

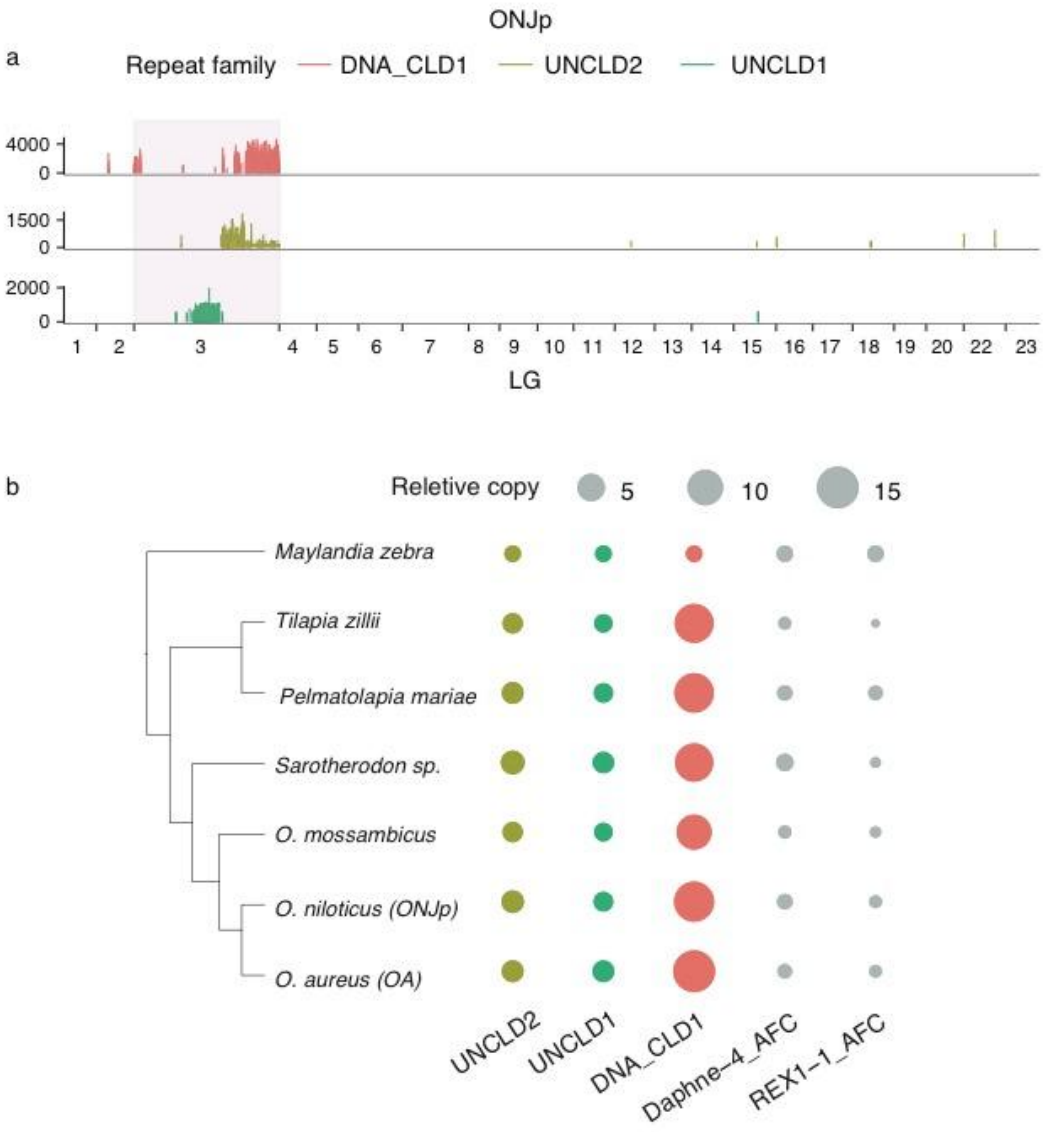


**Supplementary Figure S10 The chromosomal distribution and copy numbers of CLD repeats.** a) The density (length per 5 kb) of three transposable elements across the genome of *ONJp* that are nearly exclusively distributed on LG3. b) The copy numbers of UNCLD1, UNCLD2, DNA_CLD1 and other two family of transposable elements (grey) are compared across tilapia species and a cichlid. The relative copy numbers were calculated using the number in *M.zebra* as a baseline (the copy number in *M. zebra* became 1). Daphne-4_AFC and REX1-1_AFC were two common transposable elements randomly selected as a control.

**Supplementary Figure S11 Expression patterns of tandem gene clusters on LG3**

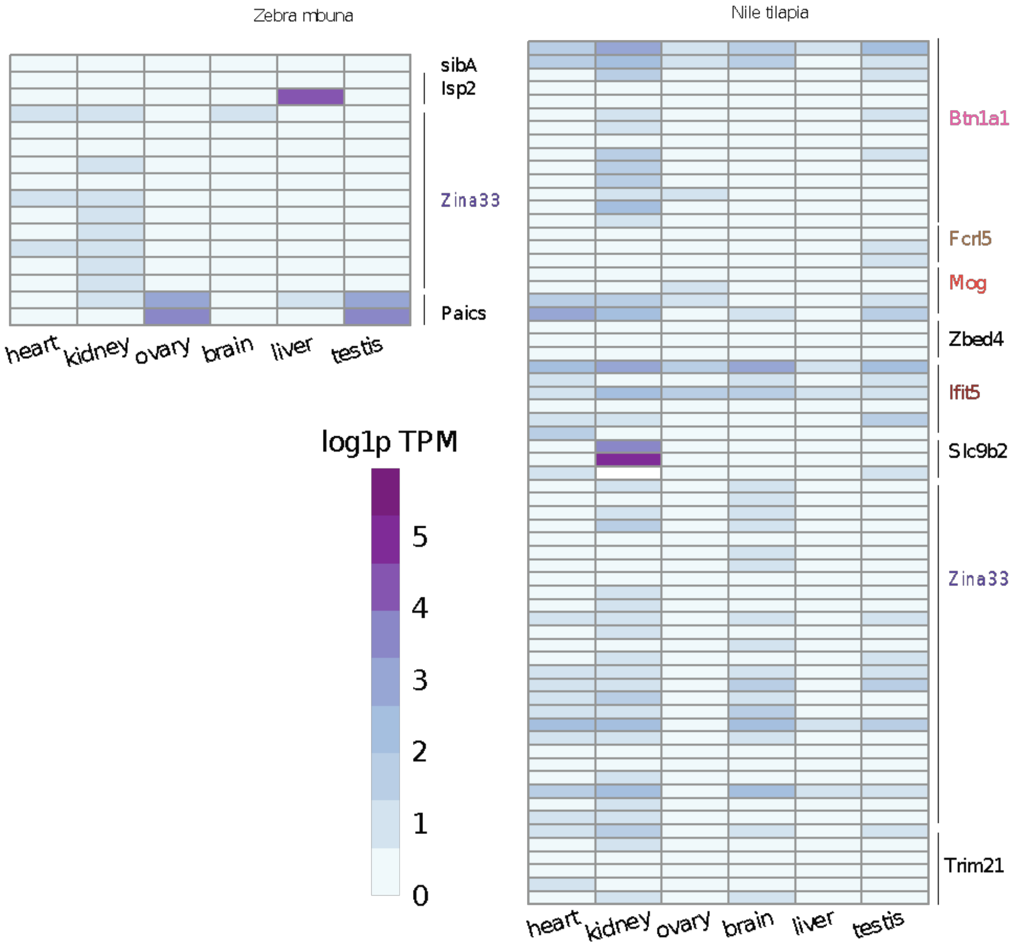


**Supplementary Figure S11 Expression patterns of tandem gene clusters on LG3.** The expression profiles of tandem gene duplicates were compared across six different tissues. The RNA-seq data were retrieved from Brawand et al. (2014)[1]. Due to limited available RNA-seq data of blue tilapia, the expression patterns were not characterized.

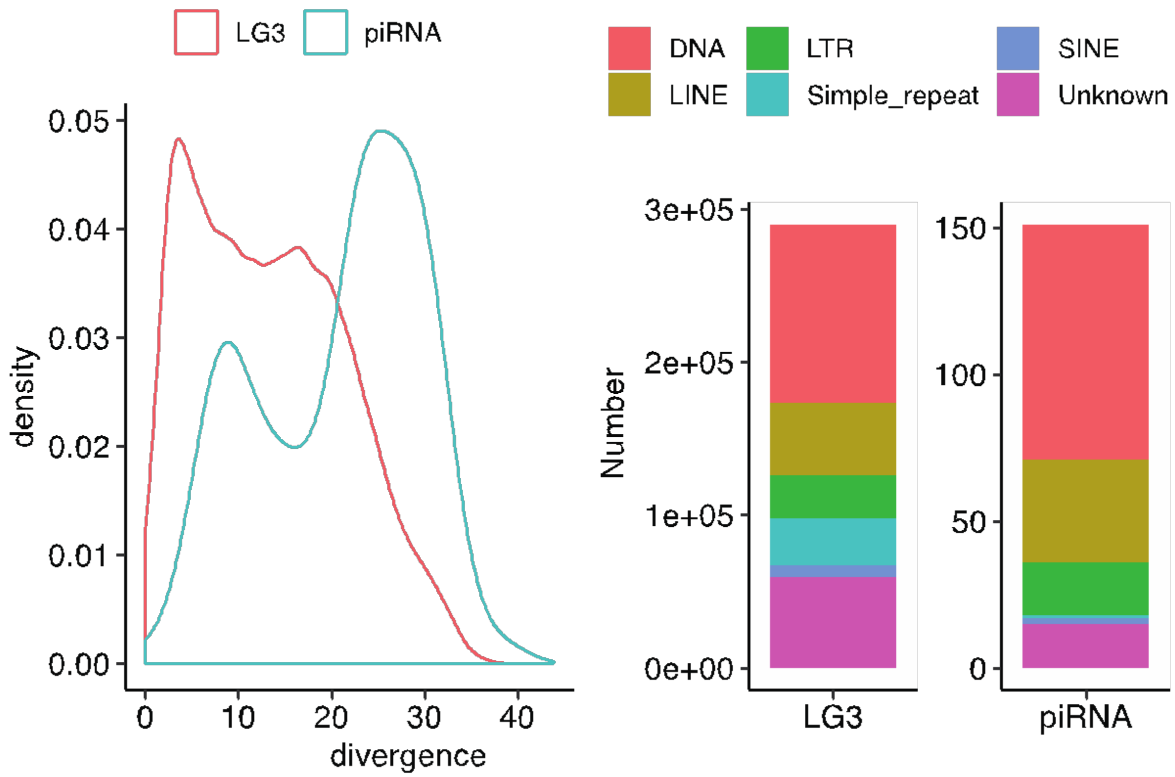


**Supplementary Figure S12 The divergence and composition of repeats targeted by piRNAs within piRNA clusters in *ONJp*.** The left panel shows the sequence divergence of repeats targeted by piRNAs derived from piRNA and that of all repeats of LG3. The right panel shows the composition of the repeats from the entire LG3 and those targeted by piRNA-cluster derived piRNAs.

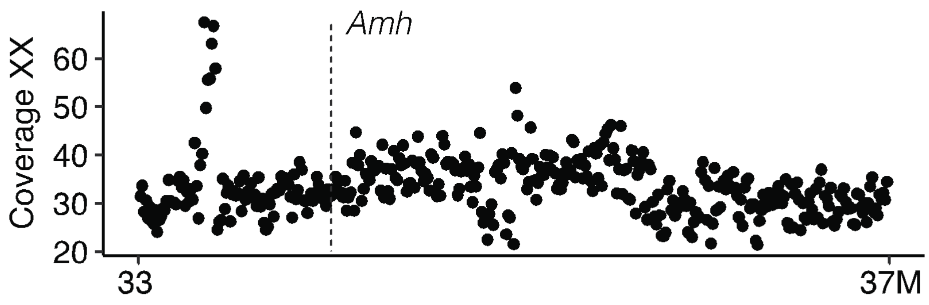


**Supplementary Figure S13 Coverage Pattern of XX reads along LG23 of *ONJp.*** The coverage of XX male was calculated for each 5k window. The location of the sex-determining gene *Amh* is indicated by a vertical dashed line. The lack of elevated coverage at the *Amh* locus was in contrast with the pattern in Figure 5b.

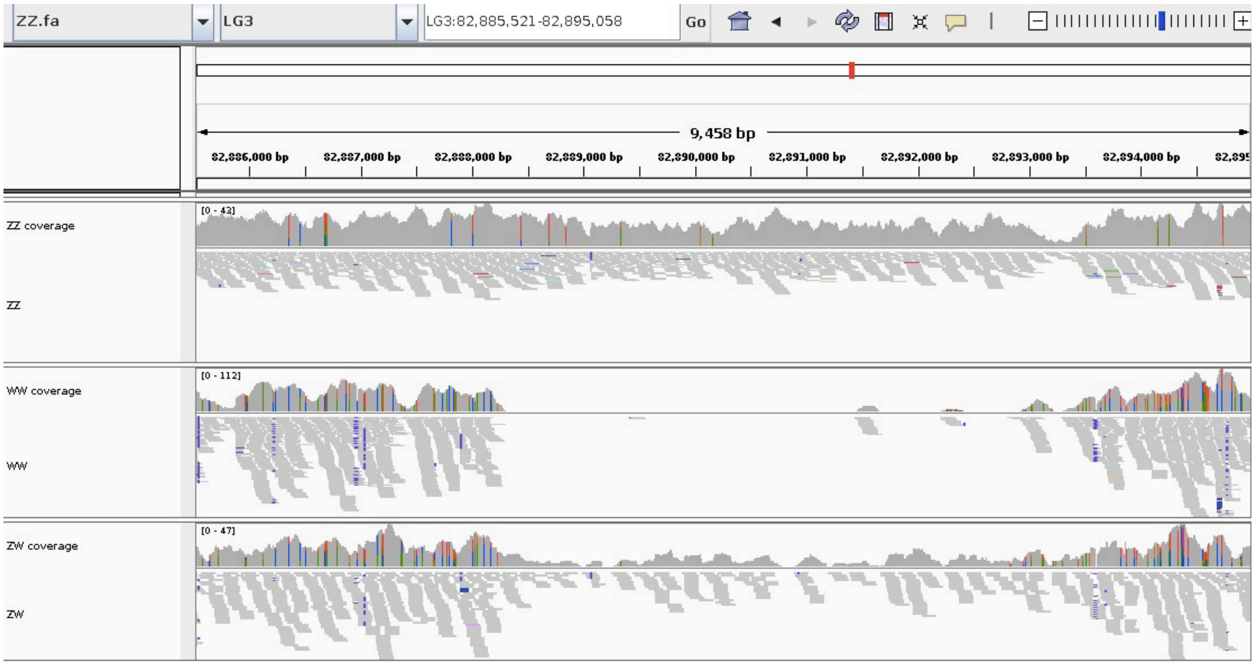


**Supplementary Figure S14 Coverage Pattern of W-linked deletion*.*** The read coverage of ZZ male, WW and ZW females are shown for the deletion region (approximately 82,888,000 - 82893,000) of Figure 5d-e. The region is not covered by any WW reads, and a reduced number of ZW reads relative to that of ZZ reads. The ZW reads also expectedly show a lack of heterozygotes at the deleted region.

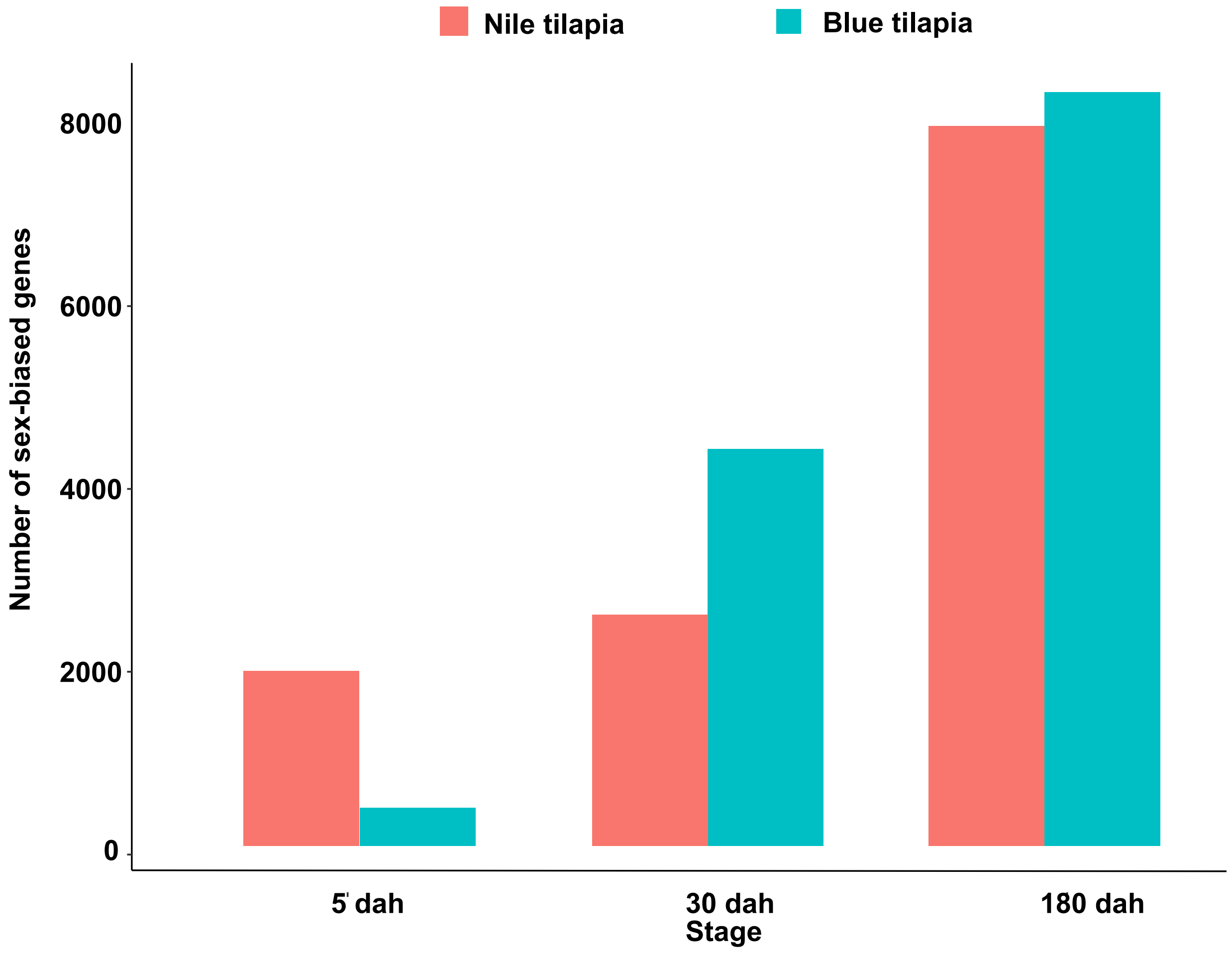


**Supplementary Figure S15 The numbers of sex-biased genes during gonadogenesis.** The number of reads for each gene was mapped and counted using Hisat2 and featureCount according to the gene model annotation. Two-fold difference in expression between female and male gonads of Nile tilapia and blue tilapia at the same same stage were used as the threshold to identify sex-biased genes.

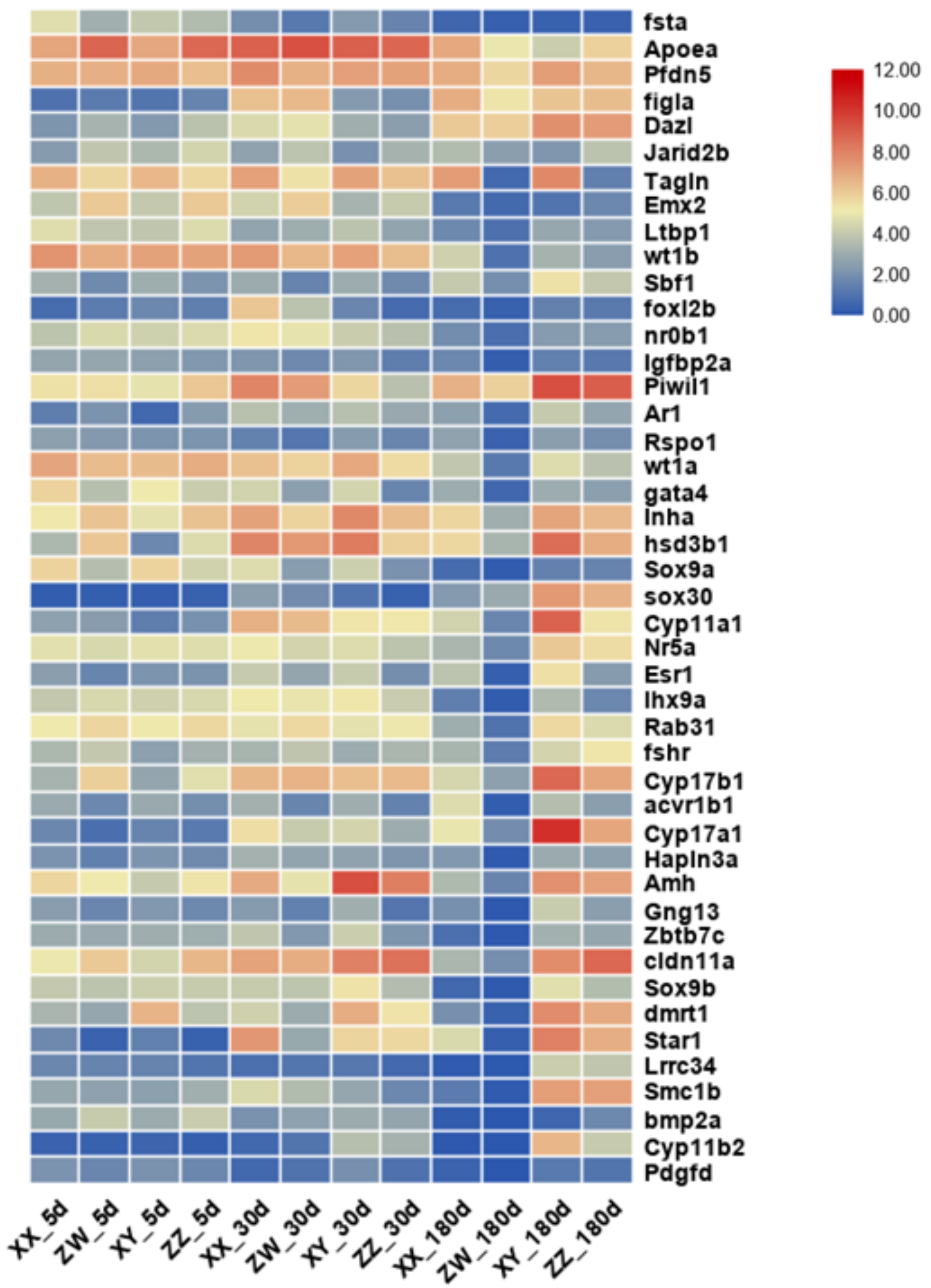


**Supplementary Figure S16 Expression patterns of known vertebrate male-determining genes. The expression of genes were calculated based on gonadal transcriptomes at 5, 30 and 180 dah. Red denotes high expression, while blue denotes low expression.**

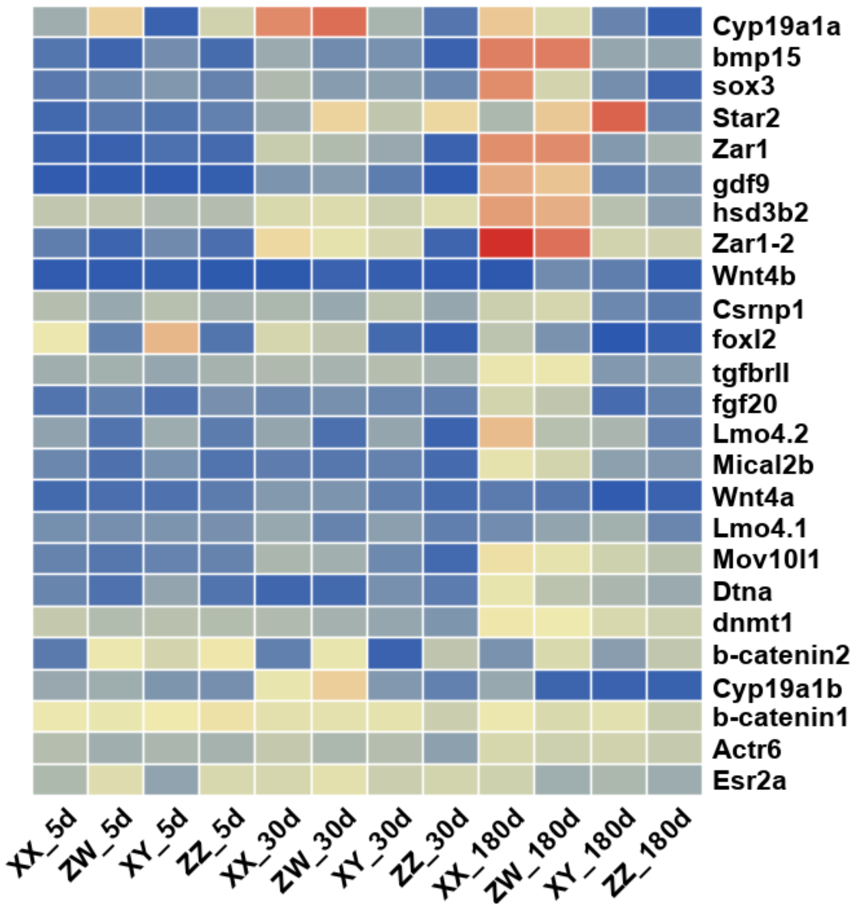


**Supplementary Figure S17 Expression patterns of known vertebrate female-determining genes. The expression of genes were calculated based on gonadal transcriptomes at 5, 30 and 180 dah. Red denotes high expression, while blue denotes low expression.**

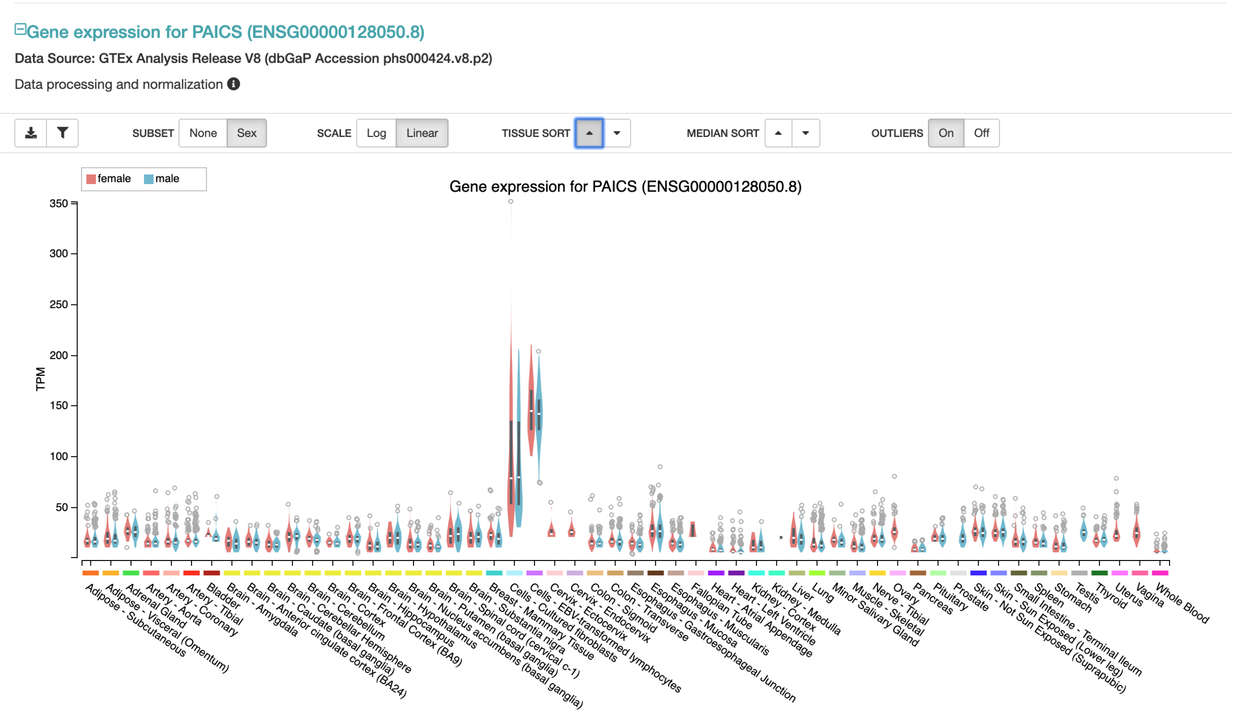


**Supplementary Figure S18 Expression patterns of human *Paics* gene across tissue types. The expression data is retrieved from GTEx Portal (https://www.gtexportal.org/home/).**

**Reference**

1. Brawand, D. *et al.* The genomic substrate for adaptive radiation in African cichlid fish. *Nature* **513,** 375–381 (2014).
